# Structural insights into regulation of the PEAK3 pseudokinase scaffold by 14-3-3

**DOI:** 10.1101/2022.09.01.506268

**Authors:** Hayarpi Torosyan, Michael D. Paul, Antoine Forget, Megan Lo, Devan Diwanji, Krzysztof Pawłowski, Nevan J. Krogan, Natalia Jura, Kliment A. Verba

**Affiliations:** Cardiovascular Research Institute, University of California San Francisco, San Francisco, CA 94158, USA; Biophysics Graduate Program, University of California, Sam Francisco, CA 94158, USA; Quantitative Biosciences Institute (QBI), University of California San Francisco, San Francisco, CA 94158, USA; Medical Scientist Training Program, University of California San Francisco, San Francisco, CA, CA 94158, USA; J. David Gladstone Institutes, San Francisco, CA 94158; Department of Cellular and Molecular Pharmacology, University of California San Francisco, San Francisco, CA 94158, USA; Department of Pharmaceutical Chemistry, University of California San Francisco, San Francisco, CA 94158, USA; Department of Molecular Biology, University of Texas Southwestern Medical Center, Dallas, TX 75390, USA; Department of Biochemistry and Microbiology, Warsaw University of Life Sciences, 02-787 Warszawa, Poland

## Abstract

The three members of the PEAK family of pseudokinases (PEAK1, PEAK2, and PEAK3) are molecular scaffolds that have recently emerged as important regulatory nodes in signaling pathways that control cell migration, morphology, and proliferation, and they are increasingly found to be mis-regulated in human cancers. While no structures of PEAK3 have been solved to date, crystal structures of the PEAK1 and PEAK2 pseudokinase domains revealed their dimeric organization. It remains unclear how dimerization plays a role in PEAK scaffolding functions, as no structures of PEAK family members in complex with their binding partners have been solved. Here, we report the cryo-EM structure of the PEAK3 pseudokinase, also adopting a dimeric state, and in complex with an endogenous 14-3-3 heterodimer purified from mammalian cells. Our structure reveals an asymmetric binding mode between PEAK3 and 14-3-3 stabilized by one pseudokinase domain and the Split HElical Dimerization (SHED) domain of the PEAK3 dimer. The binding interface is comprised of a canonical primary interaction involving two phosphorylated 14-3-3 consensus binding sites located in the N-terminal domains of the PEAK3 monomers docked in the conserved amphipathic grooves of the 14-3-3 dimer, and a unique secondary interaction between 14-3-3 and PEAK3 that has not been observed in any previous structures of 14-3-3/client complexes. Disruption of these interactions results in the relocation of PEAK3 to the nucleus and changes its cellular interactome. Lastly, we identify Protein Kinase D as the regulator of the PEAK3/14-3-3 interaction, providing a mechanism by which the diverse functions of the PEAK3 scaffold might be fine-tuned in cells.

## Introduction

By catalyzing phosphorylation, protein kinases play important roles in regulating essential cellular processes such as cell survival, proliferation, migration, and organogenesis. However, a significant subset of human kinases, termed pseudokinases, lack this vital enzymatic function due to the presence of inactivating mutations within key catalytic motifs. Instead, these catalytically inactive kinases have evolved other functionalities to propagate signaling, including allosteric modulation and scaffolding of their interaction partners^1,2^. Although pseudokinases are increasingly recognized as essential regulators of cellular signal transduction, the molecular mechanisms of most of their non-canonical kinase functions remain poorly understood.

The pseudopodium enriched atypical kinase (PEAK) family, also known as New Kinase Family 3 (NKF3), of pseudokinases is composed of three members: PEAK1 (also known as Sugen kinase [SgK] 269), PEAK2 (also known as Pragmin or SgK223), and PEAK3 (also known as chromosome 19 Open Reading Frame 35 [C19orf35]). The three PEAKs act as molecular scaffolds in pathways involved primarily in cell motility and migration^3,4^. PEAK1 overexpression increases cell motility and migration, likely due to a combination of a PEAK1-dependent increase in ERK signaling and through an ERK-independent mechanism via direct association of PEAK1 with the Src/p130Cas/Crk complex^5,6^. Notably, PEAK1 has been shown to be important for modulation of the late phase of EGF-induced EGFR signaling via its interaction with Shc1. PEAK1 binding to Shc1 results in a switch in the Shc1-associated proteome from factors involved in pro-mitogenic and pro-survival signaling to proteins involved in cytoskeletal reorganization, migration, and eventually signal termination^7^. PEAK2 was initially found to increase RhoA signaling by interacting with the Rho family GTPase Rnd2, resulting in inhibition of neurite outgrowth^8^. However, in epithelial cells, PEAK2 activity induces cell growth upon binding the C-terminal Src kinase (Csk), as a result of Src-dependent phosphorylation^9,10^. PEAK2/Csk binding increases Csk localization and activity at focal adhesions, inducing cell elongation and motility^9,10^. The emerging understanding of PEAK family signaling is that PEAKs regulate distinct cellular outcomes by selectively engaging their interaction partners.

PEAK3, the newest member of the PEAK pseudokinase family, and most distinct in its primary sequence, remained unannotated as a kinase until recently, and its functional characterization lags behind PEAK1 and PEAK2^11,12^. In fibroblast cells, PEAK3 was found to directly interact with CrkII and inhibit CrkII-dependent membrane ruffling^11,12^. Similar to what has been observed for PEAK2, the function of PEAK3 is cell type-dependent. In contrast to its effect on fibroblast cells, PEAK3 overexpression in epithelial cells was shown to increase cell motility and invasion^13^. These effects may be attributed to the multi-scaffolding capacity of PEAK3, since it interacts with a diverse set of signaling proteins, including Grb2, CrkII, Cbl, PYK2, ASAP1, and EGFR^13,14^. Reminiscent of the signal switching effect that PEAK1 has on the EGF-induced Shc1 proteome, the PEAK3 interactome is also altered by EGF stimulation, suggesting that PEAK3 has a dynamic role in signaling^13^. Importantly, in osteosarcoma and acute myeloid leukemia (AML) cell lines, overexpression of PEAK3 results in elevated AKT activity and increased cell growth^14^.

The three PEAKs have a similar domain architecture composed of an unstructured N-terminal domain followed by a pseudokinase domain^15^. The N-terminal domain contains a PEST linker (<u>P</u>roline, <u>G</u>lutamate, <u>S</u>erine, <u>T</u>hreonine-rich region), which houses numerous consensus binding motifs, including those for SH2, SH3, and PTB domains. Interestingly, the N-terminal domain of PEAK3 is approximately one-tenth the length of the N-terminal domains of PEAK1 and PEAK2 (130 residues in PEAK3 vs 1282 residues in PEAK1 and 947 residues in PEAK2).

All PEAKs are believed to utilize their N-terminal domains at least in part for scaffolding, but little is known about the mechanisms by which this happens. The pseudokinase domains in all three PEAKs have highly conserved helical regions adjoining both their N- and C-termini^11,12,16,17^. Crystal structures of truncated PEAK1 and PEAK2 lacking the unstructured N-terminal domains revealed that these pseudokinase domain-flanking helices fold into an “XL”-shaped helical bundle and facilitate dimerization by forming an “XX”-shaped high-affinity dimerization interface termed the <u>S</u>plit <u>He</u>lical <u>D</u>imerization (SHED) domain. All three PEAKs have been shown to homo- and heterodimerize in co-immunoprecipitations studies, with the pseudokinase and SHED domains being essential for these interactions^12,13,18^. The importance of this novel dimerization interface is reflected in the findings that binding of nearly all identified PEAK interaction partners is dependent on the ability of PEAK pseudokinases to dimerize^12,13,18^.

PEAKs are classified as pseudokinases due to substitutions in key catalytic motifs, including mutation of the DFG motif in PEAK1 and PEAK2 and of the HRD motif in PEAK1 and PEAK3. Crystal structures of PEAK1 and PEAK2 illustrate how these unique sequence features contribute to the lack of PEAK1 and PEAK2 catalytic activity. In these structures, the putative nucleotide-binding pockets of PEAK1 and PEAK2 are partially collapsed and occluded by bulky hydrophobic side chains, preventing nucleotide binding^11,16,17^. Due to sequence deviations in PEAK3 and the lack of PEAK3 structures, it has not been clear whether PEAK3 adopts a similar architecture. An intact DFG motif in PEAK3, as well as the presence of a conserved “catalytic lysine” with its potential salt bridge partner, a glutamate located in the predicted αC helix, presented the possibility that the nucleotide binding capability of PEAK3 might deviate from its family members.

We used cryo-electron microscopy (cryo-EM) to obtain the first structural insights into PEAK3 and the mechanisms that mediate its protein-protein interactions. Strikingly, full-length human PEAK3 purified from mammalian cells as a homodimer in complex with an endogenous 14-3-3 heterodimer, and we have obtained a 3.1 Å structure of this complex. 14-3-3s are ubiquitous regulatory proteins that act as dimeric scaffolds to modulate their client proteins through a wide array of mechanisms, such as altering protein-protein interactions, inducing conformational changes, regulating subcellular localization, and sterically blocking access to phosphorylation sites^19-22^. This has recently been exquisitely demonstrated in high resolution cryo-EM structural studies of 14-3-3s in complex with RAF and MEK kinases^23,24^. In our PEAK3 structure, key structural kinase elements, the αC helix and the activation loop, are fully resolved, in contrast to previous structures of PEAK1 and PEAK2. The N-terminal domain of PEAK3 is largely disordered, with the exception of the phosphorylated 14-3-3 binding motifs of each PEAK3 monomer that interact with each 14-3-3 monomer. Most remarkably, our structure reveals a unique secondary interface between the PEAK3 and 14-3-3 dimers that has not been previously observed in other structures of 14-3-3/client complexes. Analysis of the PEAK3/14-3-3 complex by mutagenesis demonstrates how this interaction regulates PEAK3 subcellular localization and the expansiveness of its cellular interactome.

## Results

### Structure of the PEAK3/14-3-3 complex

The long, unstructured N-terminal domains of PEAK1 and PEAK2 make it challenging to express them as recombinant full-length proteins for structural studies by X-ray crystallography^15^. We succeeded in expressing the much shorter full-length PEAK3 in mammalian Expi293F cells and purifying it for cryo-EM studies. Following affinity purification, FLAG-tagged full-length PEAK3 remained associated with a ∼ 30 kDa doublet band, which we determined to be a mixture of 14-3-3 proteins by mass spectrometry (MS) **(Figure 1A; SI Figure 1A, 1B)**. A primary sequence analysis of PEAK3 revealed the presence of a putative 14-3-3 consensus binding motif within the N-terminal domain of PEAK3 (RTQSLP, residues 66-71) (**Figure 1B, SI Figure 2A)**, which is highly conserved in evolution **(Figure 1C)**^25^.

**Figure 1:**
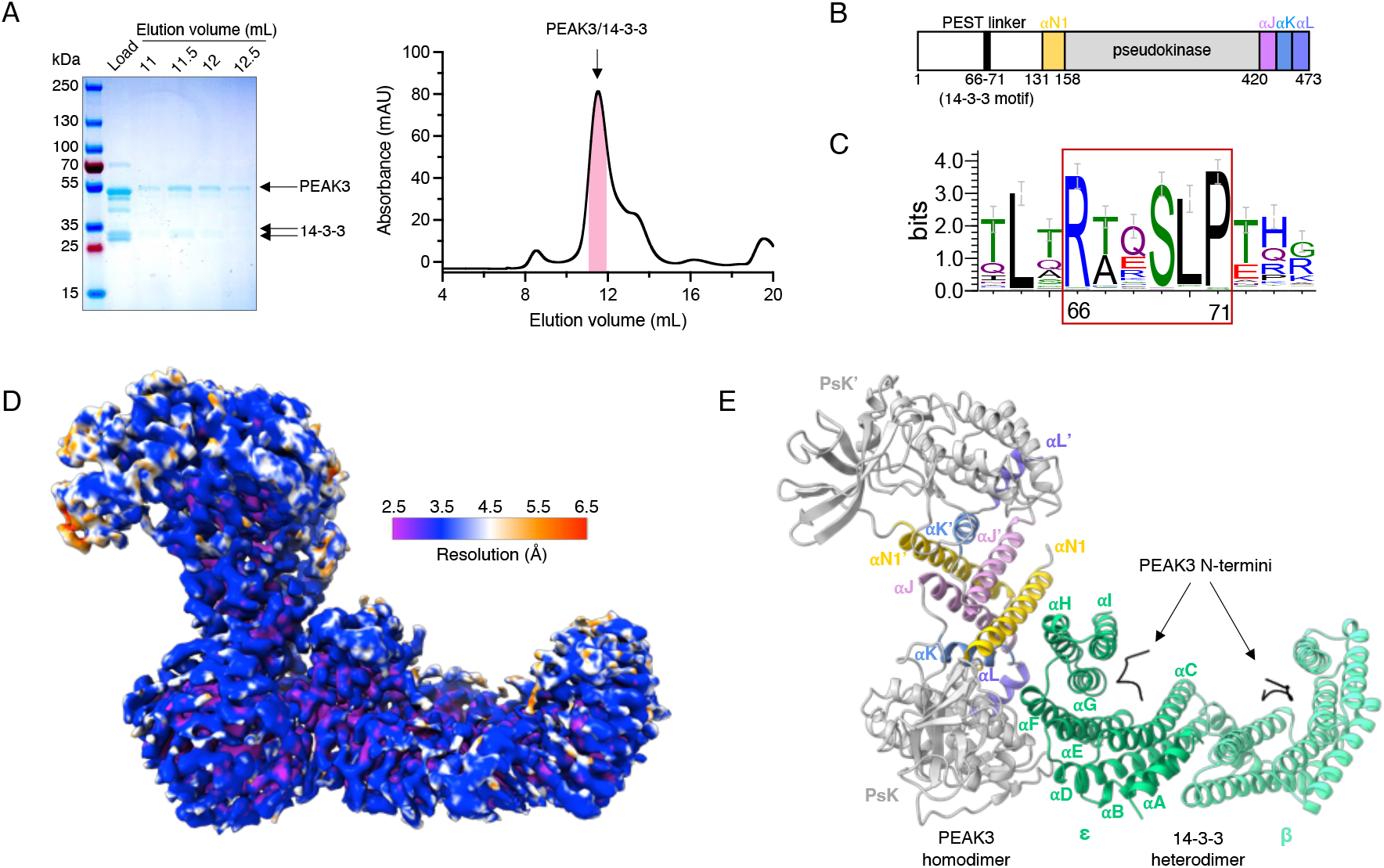
Structure of the PEAK3/14-3-3 complex. A) Representative Coomassie-stained SDS-PAGE gel analysis of the purified PEAK3/14-3-3 complex and corresponding size exclusion chromatography profile resolved on a Superdex200 10/300 Increase column. Pink bar on the chromatogram indicates collected fractions. B) Cartoon schematic of PEAK3 domain structure depicting location of the 14-3-3 binding site. C) Sequence logo demonstrating conservation of 14-3-3 binding site in PEAK3 across evolution. D) Cryo-EM map of the PEAK3/14-3-3 complex colored according to local resolution determined by ResMap. E) Structure of the PEAK3/14-3-3 complex. PEAK3 pseudokinase domains (PsK and PsK’) are in gray and the resolved regions of the N-terminal PEST linkers are shown in black. SHED domain helices αN1 are in yellow, αJ in pink, αK in blue, and αL in purple. 14-3-3ε is shown in green and 14-3-3β is shown in light green.

We observed both PEAK3/14-3-3 complex and PEAK3 homodimer particles in the cryo-EM dataset. Selective processing of the PEAK3/14-3-3 complex particles yielded a 3.1 Å structure of the PEAK3 homodimer bound to a 14-3-3 heterodimer **(Figure 1D, SI Figure 3, SI Figure 4, SI Table 1)**. Processing of PEAK3-only particles yielded a low-resolution (4.9 Å) reconstruction of the PEAK3 homodimer **(SI Figure 3, SI Figure 5A)**. This could be due to the low number of particles in the resulting reconstruction or may suggest that binding of 14-3-3 to PEAK3 stabilizes the otherwise dynamic PEAK3 homodimers. Comparison of the PEAK3/14-3-3 structure with the low-resolution structure of the PEAK3 homodimer alone did not reveal major differences within the PEAK3 dimer, with an overall average RMSD of 0.623 Å **(SI Figure 5B)**.

Previously, PEAK1 and PEAK2 pseudokinase dimers have been shown to organize into higher-order oligomers^17,18^. Concordant with this, a subset of 2D class averages in our PEAK3/14-3-3 cryo-EM dataset was also consistent with higher-order PEAK3 oligomers, both in the absence and presence of 14-3-3. However, the particle counts for these complexes were too low to enable 3D reconstructions **(SI Figure 6A, B)**. To investigate if such oligomeric complexes of PEAK3 could form using the same interface observed in PEAK2, and presumably in PEAK1, even in the presence of 14-3-3 binding, we built structural models of PEAK3 oligomers using the interface observed in the crystal structure of PEAK2^17^. This analysis revealed that the PEAK3 dimer is able to engage both 14-3-3 and another PEAK3 dimer simultaneously via the same monomer or different monomers without steric clashes **(SI Figure 6C)**.

MS analysis of the purified PEAK3/14-3-3 complex used for cryo-EM identified six of the seven isoforms of human 14-3-3 (β, γ, ε, η, θ, and ζ) **(SI Figure 1A, 1B)** and phosphorylation of the Ser 69 residue within the 14-3-3 binding motif of PEAK3 **(SI Figure 1C**). In the cryo-EM map, the phosphorylated Ser 69 was clearly identifiable in both 14-3-3 binding grooves, allowing us to manually build the neighboring peptide sequences (RTQS^69^LP), presumably coming from both N-terminal domains of the PEAK3 dimer. In the structure, the PEAK3 homodimer is docked to one side of the 14-3-3 dimer cradle, with only one monomer of PEAK3 forming an extensive interface with one of the 14-3-3 monomers, yielding a buried surface area (BSA) of ∼1453 Å^2^ **(Figure 1E)**. Based on the cryo-EM map, we were able to unambiguously identify this 14-3-3 monomer as the ε isoform (**SI Figure 7A-7G)**, while the distal 14-3-3 monomer was most consistent with the β isoform **(SI Figure 7H-7N)**. Because MS analysis identified multiple 14-3-3 isoforms, all of which exhibit high sequence conservation, we cannot rule out a mixture of 14-3-3 isoforms in the β monomer density. Nevertheless, our final model was constructed with the ε and β isoforms of 14-3-3, which most closely match the cryo-EM density. Although it has been well appreciated that 14-3-3ε preferentially heterodimerizes with other 14-3-3 isoforms rather than forming homodimers, likely due to the formation of an additional salt-bridge at the dimerization interface^26^, our structure provides the first high resolution experimental view of such a heterodimer. With the exception of the 14-3-3 binding motifs, discussed later, the remainder of the PEAK3 N-terminal domain of each monomer was not resolved, consistent with its predicted intrinsic flexibility **(Figure 1E)**^15^.

### SHED-mediated dimerization is conserved in PEAK3

Similar to the previously seen dimerization poses of PEAK1 and PEAK2, in our structure, PEAK3 dimerizes via the “XL”-shaped helical bundles flanking each pseudokinase domain. Consistent with previous predictions based on sequence conservation and homology modeling^12^, these individual “XL” helical bundles come together to form the canonical “XX”-shaped SHED domain burying a total surface area of 2726 Å^2^ **(Figure 2A)**. The “XL”-shaped helical bundle within each PEAK3 monomer is held together by a tightly packed network of hydrophobic interactions involving the αN1, αJ, αK, and αL helices, largely conserved in PEAK1 and PEAK2 **(Figure 2B)**. Notably, Leu 146 of the αN1 and Cys 453 of the αK helix contribute to this network. When individually mutated, these residues are sufficient to disrupt PEAK3 homo- and heterodimerization^12,13^ and ablate its interaction with almost all identified binding partners^13,14^. The SHED domain is stabilized primarily via hydrophobic interactions between the αN1 and αJ helices of the two PEAK3 monomers **(Figure 2C)**. This interface contains Ala 436 on the αJ helix, mutation of which has also been shown to disrupt PEAK3 dimerization and its scaffolding functions^12,13^.

**Figure 2:**
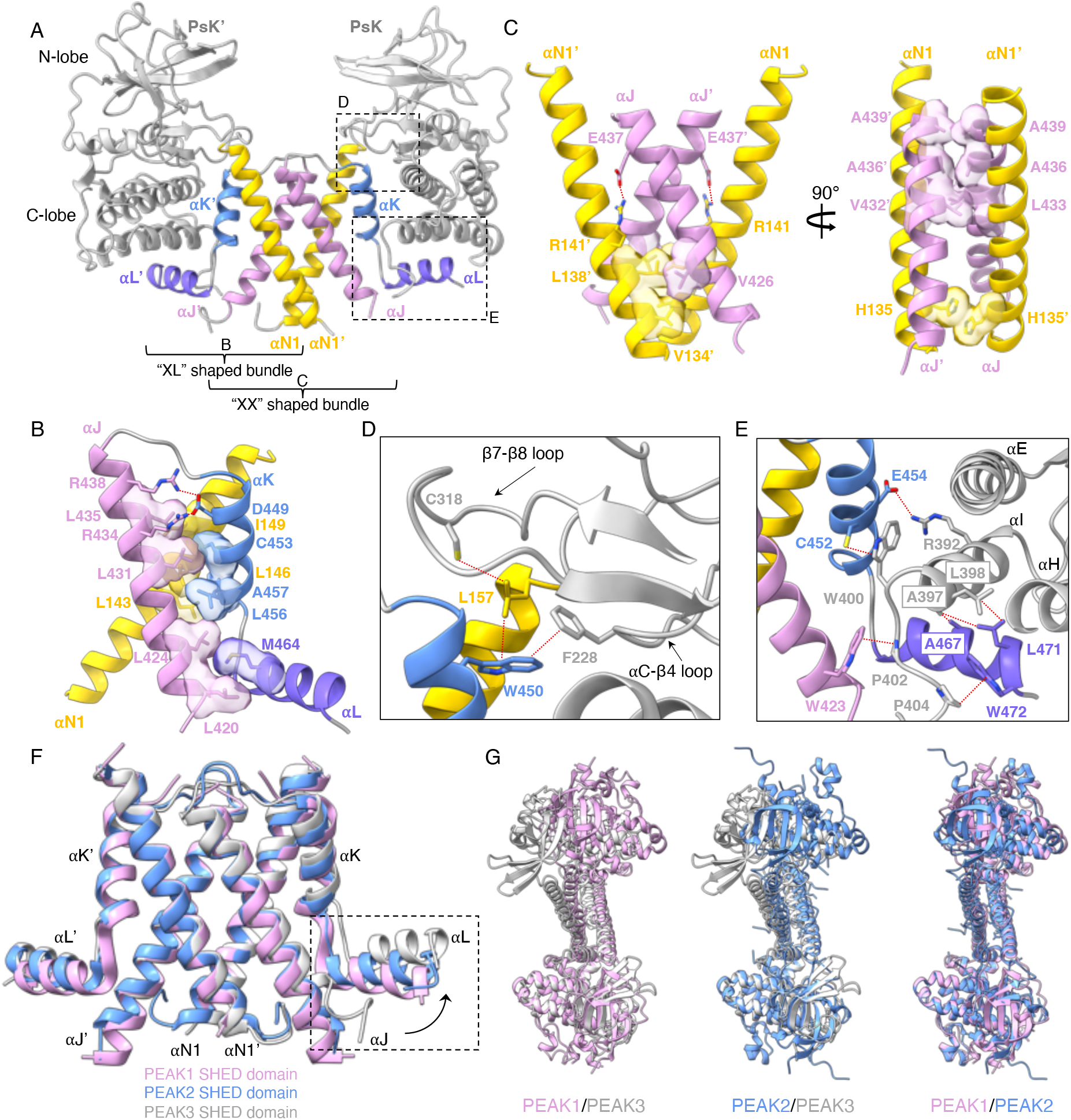
SHED-dependent dimerization of PEAK3. A) Overall structure of the PEAK3 dimer as part of the PEAK3/14-3-3 complex (14-3-3 is not shown). B) XL shaped helical bundle of one PEAK3 monomer, highlighting intramolecular interactions. C) Intermolecular interactions between helices αN1 and αJ of each PEAK3 monomer within the SHED dimer. D-E) Zoomed-in views of the interactions between the SHED domain and pseudokinase/N-lobe (D) and pseudokinase/C-lobe (E). F) Structural alignment of PEAK family SHED domains (PDB ID: 6BHC for PEAK1, and PDB ID: 5VE6 for PEAK2) along αN1 and αJ helices, depicting movement of αJ and αL helices in PEAK3 relative to their position in PEAK1 and PEAK2. G) Overlay of the PEAK1 (PDB ID: 6BHC), PEAK2 (PDB ID: 5VE6) and PEAK3 dimer structures aligned along one pseudokinase domain (shown at the bottom of each overlay). Dashed lines indicate interactions with distances ≤ 4 Å.

Like in PEAK1 and PEAK2, SHED domain helices not only participate in the dimerization interface, but they also make extensive interactions with the pseudokinase domain of PEAK3 (**Figure 2D, 2E**). The N-lobe of the pseudokinase domain, specifically the αC-β4 loop Phe 228 (PEAK1: Phe 1385, PEAK2: Phe 1045), forms a hydrophobic contact with Trp 450 (PEAK1: Trp 1722, PEAK2: Trp 1382) of the αK helix. Trp 450 additionally links the αN1 helix to the β7-β8 loop of the N-lobe by positioning Leu 157 (PEAK1: Phe 1309, PEAK2: Phe 974) to interact with Cys 318 (PEAK1: unresolved, PEAK2: Cys 1156) **(Figure 2D)**. The C-lobe of the pseudokinase domain forms a more substantial interface with the αJ, αK and αL helices of the SHED domain.

In particular, the αK residues, Cys 452 (PEAK1: Cys 1724, PEAK2: Cys 1384) and Glu 454 (PEAK1: Gln 1726, PEAK2: Gln 1386), form hydrophobic and electrostatic contacts with Trp 400 (PEAK1: Trp 1668, PEAK2: Trp 1340) and Arg 392 (PEAK1: Lys 1660, PEAK2: Lys 1322) of the αI helix, respectively. In turn, Leu 398 (PEAK1: Leu 1666, PEAK2: Leu 1328) and Ala 397 (PEAK1: Cys 1665, PEAK2: Cys 1327) of the αI helix pack against Ala 471 (PEAK1: Leu 1743, PEAK2: Leu 1403) and Leu 467 (PEAK1: Ile 1739, PEAK2: Ser 1399) of the αL helix, while Pro 402 (PEAK1: Pro 1670, PEAK2: Pro 1332), located on the C-terminus of the pseudokinase domain, forms an additional hydrophobic contact with Trp 423 (PEAK1: Trp 1694, PEAK2: Trp 1354) of the αJ helix **(Figure 2E)**. The SHED domain in PEAK3 has the greatest rotation angle among all PEAK structures, and this rotation is caused by an outward movement of the αJ helix, which in turn forces the αL helix to rotate upwards **(Figure 2F)**. Consequently, the pseudokinase domain of PEAK3, which interfaces with the αL helix via its C-lobe, is pushed upward and to the back relative to PEAK1 and PEAK2, resulting in a more twisted overall architecture in the PEAK3 homodimer **(Figure 2G)**.

### Asymmetric binding mode of the PEAK3/14-3-3 complex

Binding of 14-3-3s to client proteins is typically mediated by two sets of contacts, termed the primary and secondary interactions^26^. The primary interaction involves binding of the client to the conserved amphipathic groove formed by the αI, αE, αG, and αC helices of 14-3-3 via a consensus 14-3-3 binding motif which contains a phosphorylated serine or threonine residue^26^. The secondary interaction involves a larger interface between the globular domain of the client protein and 14-3-3^26^. Although the disordered N-terminal domain of PEAK3 is largely unresolved in our structure (residues 1-127), the cryo-EM map does contain robust density for the 14-3-3 primary interaction motif (residues 66-71), with clear density for the phosphate group of Ser 69 **(Figure 3A)**. The Ser 69 phosphate group forms contacts with the positively charged groove of 14-3-3, including the canonical Arg (ε-57/β-58), Arg (ε-130/β-129), Tyr (ε-131/β-130) triad as well as a Lys (ε-50/β-51) within the αE helix **(Figure 3B)**. The PEAK3/14-3-3 interaction maintains hydrogen bonding between the conserved residues of 14-3-3 (ε-Asn 175/β-Asn 176 and ε-Asn 227/β-Asn 126) and Leu 70 of PEAK3 and the backbone of PEAK3’s consensus peptide, respectively **(Figure 3B)**.

**Figure 3:**
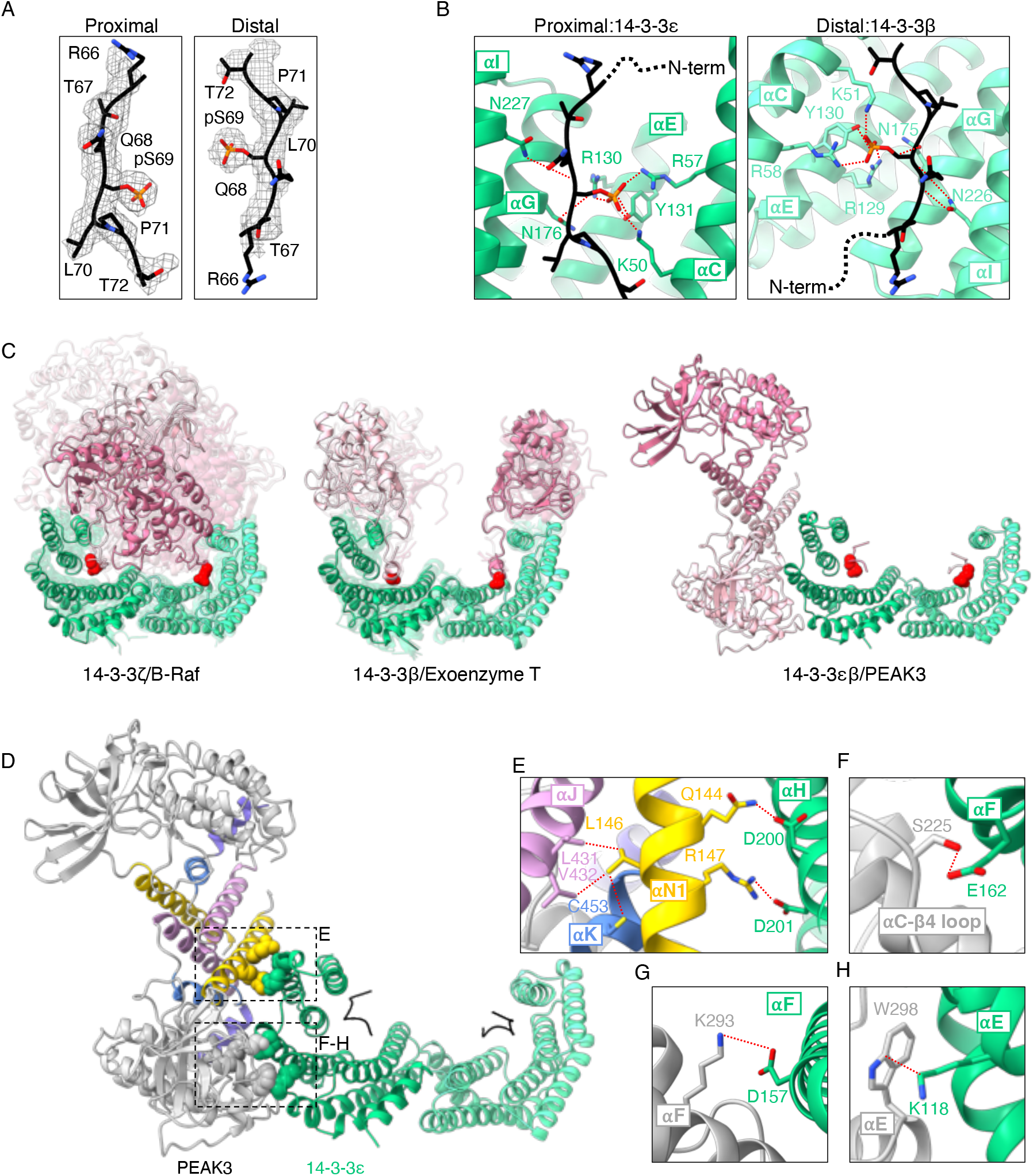
Asymmetric binding mode of the PEAK3/14-3-3 complex. A) Zoomed-in view of PEAK3 N-terminal 14-3-3 consensus binding sites superimposed with the cryo-EM map, demonstrating occupancy of both canonical substrate binding grooves in 14-3-3 by phosphorylated 14-3-3 consensus sequences of PEAK3. B) PEAK3 binding to the amphipathic grooves of 14-3-3ε and 14-3-3β. C) Known binding modes of 14-3-3/substrate complexes found in the Protein Data Bank. 14-3-3 monomers are shown in green and light green, 14-3-3 substrate monomers are shown in pink and light pink, and phosphorylated residues or phosphomimetics which engage the 14-3-3 binding grooves are shown as red spheres. See Materials and Methods for alignment details. D) PEAK3/14-3-3 structure highlighting PEAK3 SHED domain (box E) and pseudokinase domain (box F-H) interactions with 14-3-3ε. E-H) Zoomed-in view of PEAK3/14-3-3ε secondary interface, highlighting key interactions (dashed lines indicate interactions with distances ≤ 4 Å).

The secondary interaction between PEAK3 and 14-3-3 in our model is quite unusual because it is located outside of the 14-3-3 cradle. Only one monomer of 14-3-3 engages the PEAK3 dimer at this interface via helices αH, αE, and αF of 14-3-3ε **(Figure 1E)**. This binding mode is distinct from any other 14-3-3/client complexes found in the Protein Data Bank (PDB), which can be sorted into two separate categories. The first has the client protein sitting within the cradle formed by the 14-3-3 dimer, interacting with the αI helix or both αI and αH helices of 14-3-3, as observed with the B-Raf/14-3-3 complex **(Figure 3C: left)**^23^. A second binding mode has the globular domains of the client sitting atop the sides of the 14-3-3 cradle, forming contacts with both αI and αH helices, as seen in the Exoenzyme T/14-3-3 complex **(Figure 3C: middle)**^27^. To our knowledge, 14-3-3/client complexes where the dimeric client sits outside of the 14-3-3 cradle and the secondary interface only forms contacts with one 14-3-3 monomer, as observed in the PEAK3/14-3-3 complex **(Figure 3C: right)**, have never been captured in crystal structures.

The unique secondary interface in the PEAK3/14-3-3 complex is stabilized by several polar contacts between the SHED and the pseudokinase domain of PEAK3 and 14-3-3ε **(Figure 3D)**. Residues Gln 144 and Arg 147 of the αN1 helix within the SHED domain of PEAK3 form a hydrogen bond and a salt-bridge with Asp 200 and Asp 201 belonging to the 14-3-3ε αH helix, respectively **(Figure 3E)**. Leu 146, a residue we and others have previously demonstrated to be important for PEAK3 dimerization^12,13^, sits between Gln 144 and Arg 147, but faces inwards towards the SHED domain interface. Leu 146 makes hydrophobic contacts with Leu 431 and Val 432 of the αJ helix and Cys 453 of the αK helix, another residue involved in PEAK3 dimerization **(Figure 3E)**^12,13^. The close proximity of Gln 144 and Arg 147 to Leu 146 suggests that PEAK3 dimerization likely results in the proper positioning of Gln 144 and Arg 147 to interact with 14-3-3, increasing the affinity of the PEAK3/14-3-3 interaction. Several residues within the pseudokinase domain of PEAK3 also make polar contacts with 14-3-3ε. These include Ser 225 of the β3-β4 loop **(Figure 3F)**, which hydrogen bonds with Glu 162 of the αF helix of 14-3-3ε, as well as Lys 293 and Trp 298 **(Figure 3G, 3H)**, which sit on the αE helix of PEAK3 and interact with the αF helix Asp 157 and the αE helix Lys 118 of 14-3-3ε, respectively. This large interface contributed by the pseudokinase domain of PEAK3 likely stabilizes the unusual asymmetric interaction with 14-3-3ε.

### Unique features of the PEAK3 pseudokinase domain

Our structure shows that the PEAK3 pseudokinase domain adopts the canonical kinase fold, composed of a β-strand rich N-lobe and an α-helical C-lobe, although it lacks the αG helix— a feature that distinguishes it from PEAK1 and PEAK2 **(Figure 4A, 4B)**. There are several other key structural elements that are unique to the PEAK3 pseudokinase compared to PEAK1 and PEAK2. For example, the activation loop, which is largely disordered in structures of PEAK1 and PEAK2, is fully resolved in PEAK3 and adopts an extended conformation. Notably, PEAK3’s activation loop is approximately 2/3 the length of that of PEAK1 and PEAK2 **(Figure 4A)**, which could explain its increased stability and thus resolvability. The DFG motif, only present in PEAK3, is in the “DFG-in” state in our structure, as seen in the canonical “active” conformation in kinases **(Figure 4C)**. Additionally, the αC helix is fully resolved in our PEAK3 structure, in contrast to other PEAK structures, and it adopts a conformation similar to the αC helix in active PKA **(Figure 4D)**. The partially resolved αC helix in PEAK1 is in a similar conformation, while the αC helix is completely unresolved in the PEAK2 structure **(Figure 4B)**. Interestingly, sequence alignment of the αC helix region of all three PEAKs shows that the C-terminal portion, which forms an ordered helix in the PEAK1 and PEAK3 structures, is absent in PEAK2 **(Figure 4A)**. Our ability to fully resolve the αC helix and the activation loop of PEAK3 is likely also due to stabilization of these regions upon 14-3-3 binding. In our structure, both the αC helix and the activation loop pack against the 14-3-3ε monomer, contributing to the already extensive interface formed between the PEAK3 and 14-3-3 dimers **(Figure 4E)**. Notably, the region of the PEAK3 activation loop which interfaces with 14-3-3 is rich in proline and glycine residues (residues 338-347: “PPGPPGSPGP”) and is not present in PEAK1 or PEAK2.

**Figure 4:**
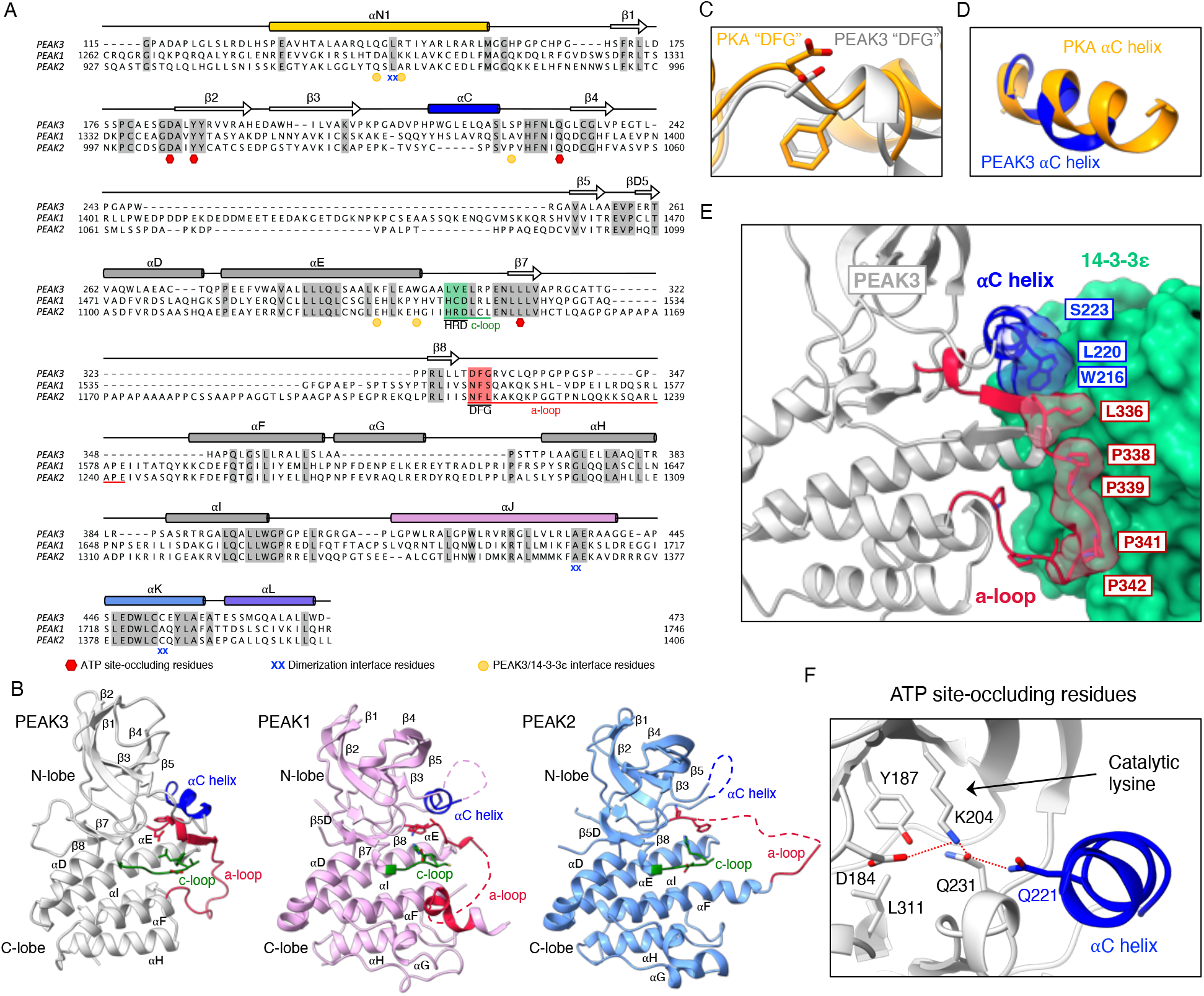
Unique features of the PEAK3 pseudokinase domain. A) Protein sequence alignment of PEAK1, PEAK2, and PEAK3 depicting secondary structure elements, sequence motifs corresponding to canonical catalytic motifs in active kinases, and key dimerization interface residues based on structures of PEAK1 (PDB ID: 6BHC), PEAK2 (PDB ID: 5VE6) and PEAK3 (PEAK3/14-3-3 complex). Conserved residues are shaded in gray, αC helix is shown in blue while catalytic and activation loops with their corresponding motifs are highlighted in green and red, respectively. Residues involved in PEAK3 dimerization and those shown to occlude the pseudoactive site and PEAK3/14-3-3 secondary interface residues are marked by the indicated symbols. B) Comparison of PEAK3 pseudokinase domain (light gray) with PEAK1 (light pink, PDB ID: 6BHC) and PEAK2 (light blue, PDB ID: 5VE6) pseudokinase domains with key structural elements highlighted (catalytic loop (c-loop) in green, activation loop (a-loop) in red, αC helix in blue). Zoomed-in view of the C) DFG motif and D) αC helix in PEAK3, overlayed on the structure of PKA (PDB ID: 1ATP), aligned along entire kinase domain. E) Zoomed-in view of the packing between αC helix and activation loop of PEAK3 and the 14-3-3ε monomer. F) Zoomed-in view of the pseudoactive site of PEAK3 showing the occluded nucleotide-binding site, the orientation of the catalytic lysine and the substituted Gln 221 in the αC helix.

Despite an active-like conformation of helix αC in PEAK3 and the presence of the “catalytic lysine,” the essential salt bridge which forms between the catalytic lysine and the αC helix glutamate in active kinases is absent in PEAK3, as the αC helix glutamate in PEAK3 is substituted with Gln 221 **(Figure 4F)**. Additionally, like in the structures of the PEAK1 and PEAK2 pseudokinases, the nucleotide-binding site in PEAK3 is sterically occluded, preventing nucleotide binding. In PEAK3, the side chains of Asp 184, Tyr 187, Gln 231, and Leu 311 occupy the pocket, with Asp 184 and Gln 231 “hijacking” PEAK3’s “catalytic lysine” (Lys 204), while Gln 221 of the αC helix hydrogen bonds with Gln 231 **(Figure 4F)**. Therefore, although PEAK3 mimics some organizational features associated with active conformation of protein kinases, the fully occluded active site we observe is consistent with its classification as a pseudokinase.

### The PEAK3/14-3-3 interaction is dependent on a canonical 14-3-3 binding motif and is further stabilized by the secondary interface

We tested the contribution of the individual binding interfaces in the PEAK3/14-3-3 structure to complex formation via co-immunoprecipitation of endogenous 14-3-3 with transiently expressed FLAG-tagged PEAK3 variants in HEK293 cells **(Figure 5A)**. Endogenous 14-3-3 robustly co-immunoprecipitated with wild type (WT) PEAK3, while mutation of the consensus site phosphorylated serine (S69A) was sufficient to completely abolish this interaction **(Figure 5B)**. Next, we examined whether the secondary interface between PEAK3 and 14-3-3ε, involving PEAK3’s SHED and pseudokinase domains, was important for the interaction. We generated mutants of PEAK3 aimed at breaking critical interactions between the αN1 helix of the SHED domain (R147A) or the pseudokinase domain (S225A, K293A, W298A, K293A/W298A) and 14-3-3ε. Mutations in the secondary interface also diminished co-immunoprecipitation of PEAK3 with endogenous 14-3-3, with the disruption of the SHED domain interaction involving the αN1 helix (R147A) leading to an approximately 75% reduction in 14-3-3 association **(Figure 5C, 5D)**. Most of the mutations within the pseudokinase domain (K293A, W298A, K293A/W298A) resulted in a more modest, but statistically significant, decreases in 14-3-3 association **(Figure 5C, 5D)**. However, none of the mutations designed to break the secondary interface were able to completely ablate the PEAK3/14-3-3 interaction, suggesting that this interface likely serves to stabilize the complex after engagement of the phosphorylated 14-3-3 consensus sequence.

**Figure 5:**
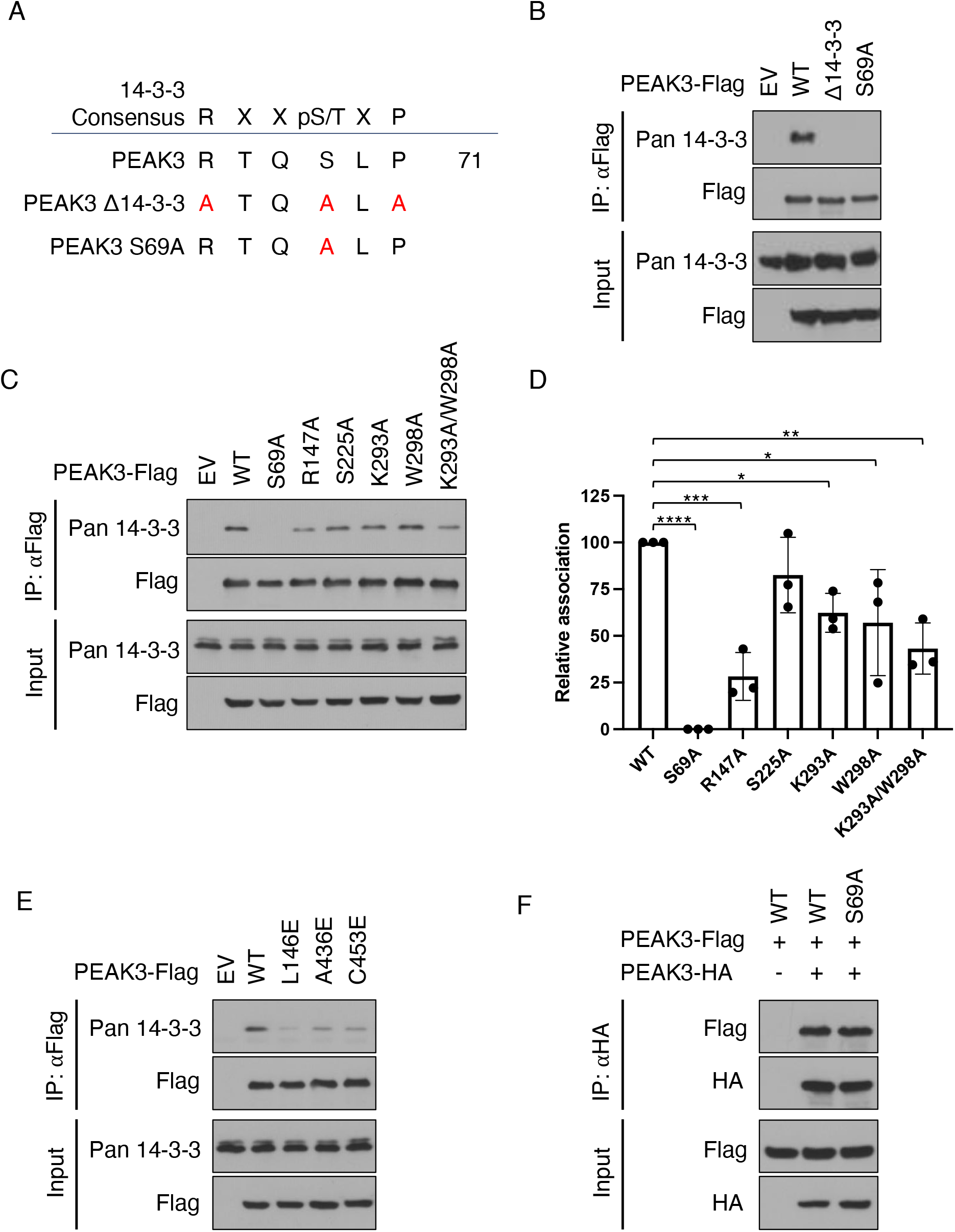
PEAK3 homodimerization and both 14-3-3 binding interfaces are critical for PEAK3/14-3-3 complex formation. A) Diagram of PEAK3 mutants generated to probe PEAK3 binding to the primary site on 14-3-3. B) Co-immunoprecipitation of the endogenous 14-3-3 with FLAG-tagged WT PEAK3 and PEAK3 mutants (Δ14-3-3, S69A) transiently expressed in HEK293 cells. C) Representative co-immunoprecipitation of the endogenous 14-3-3 with PEAK3 variants carrying mutations in the secondary 14-3-3 binding interface (R147A, S225A, K293A, W298A, K293A/W298A) transiently expressed in HEK293 cells. D) Quantification of co-immunoprecipitation data shown in panel C plotted as the mean with standard deviation from 3 independent experiments, *p < 0.05. E) Co-immunoprecipitation of the endogenous 14-3-3 with the homodimerization-deficient PEAK3 mutants (L146E, A436E, C453E) transiently expressed in HEK293 cells. F) Co-immunoprecipitation of FLAG-tagged and HA-tagged wild-type PEAK3 and PEAK3 S69A transiently expressed in HEK293 cells. In panels B-F, protein levels were detected with the indicated antibodies by Western Blot. All co-immunoprecipitation data are representative of at least 3 independent experiments.

### PEAK3 homodimerization is necessary for high-affinity binding to 14-3-3

We and others have previously shown that PEAK3 homodimerization is essential for its ability to interact with its binding partners and for its signaling^12,13^. Therefore, we asked whether PEAK3 dimerization is also necessary for 14-3-3 binding. Dimerization-deficient mutants of PEAK3 (L146E, A436E, C453E)^12^ were significantly impaired in binding the endogenous 14-3-3, as measured by co-immunoprecipitation, emphasizing the importance of PEAK3 dimerization for a stable 14-3-3 interaction **(Figure 5E)**. We then looked if the reciprocal is true, as one of the well-characterized functions of 14-3-3 proteins is mediating protein-protein interactions between client proteins, including dimerization^28,29^. We thus tested whether 14-3-3s contribute to PEAK3 dimerization, by co-expressing HA-tagged and FLAG-tagged WT or S69A variants of PEAK3 in HEK293 cells and measuring their ability to interact by co-immunoprecipitation. The FLAG-tagged S69A PEAK3 mutant interacted with the HA-tagged S69A PEAK3 to the same extent as WT PEAK3 **(Figure 5F)**, indicating that 14-3-3 binding is not necessary for PEAK3 dimerization.

### 14-3-3 regulates intracellular localization of PEAK3

A well-documented functional role of 14-3-3 scaffolds is the regulation of cellular localization of their client proteins, for example, shuttling client proteins between the cytosol and the nucleus^30^. To test if 14-3-3 binding regulates PEAK3 subcellular localization, we used immunofluorescence-based imaging of FLAG-tagged WT PEAK3 and 14-3-3 binding deficient mutant (S69A). We found that WT PEAK3 uniformly distributed throughout the cytoplasm of COS-7 cells as previously observed^12^; in contrast, the PEAK3 S69A mutant was predominantly localized in the nucleus of COS-7 cells **(Figure 6A)**. Mutations that disrupted the secondary 14-3-3 interface also increased PEAK3 nuclear localization, albeit to a smaller extent **(Figure 6B)**, consistent with their weaker effect on disrupting PEAK3/14-3-3 binding **(Figure 5D)**. These data demonstrate that 14-3-3 has a profound effect on PEAK3 cellular localization by limiting its nuclear localization upon binding. While 14-3-3s typically alter client nuclear localization by obscuring nuclear import or export signals^21,31,32^, we have not identified such signals in the PEAK3 sequence.

**Figure 6:**
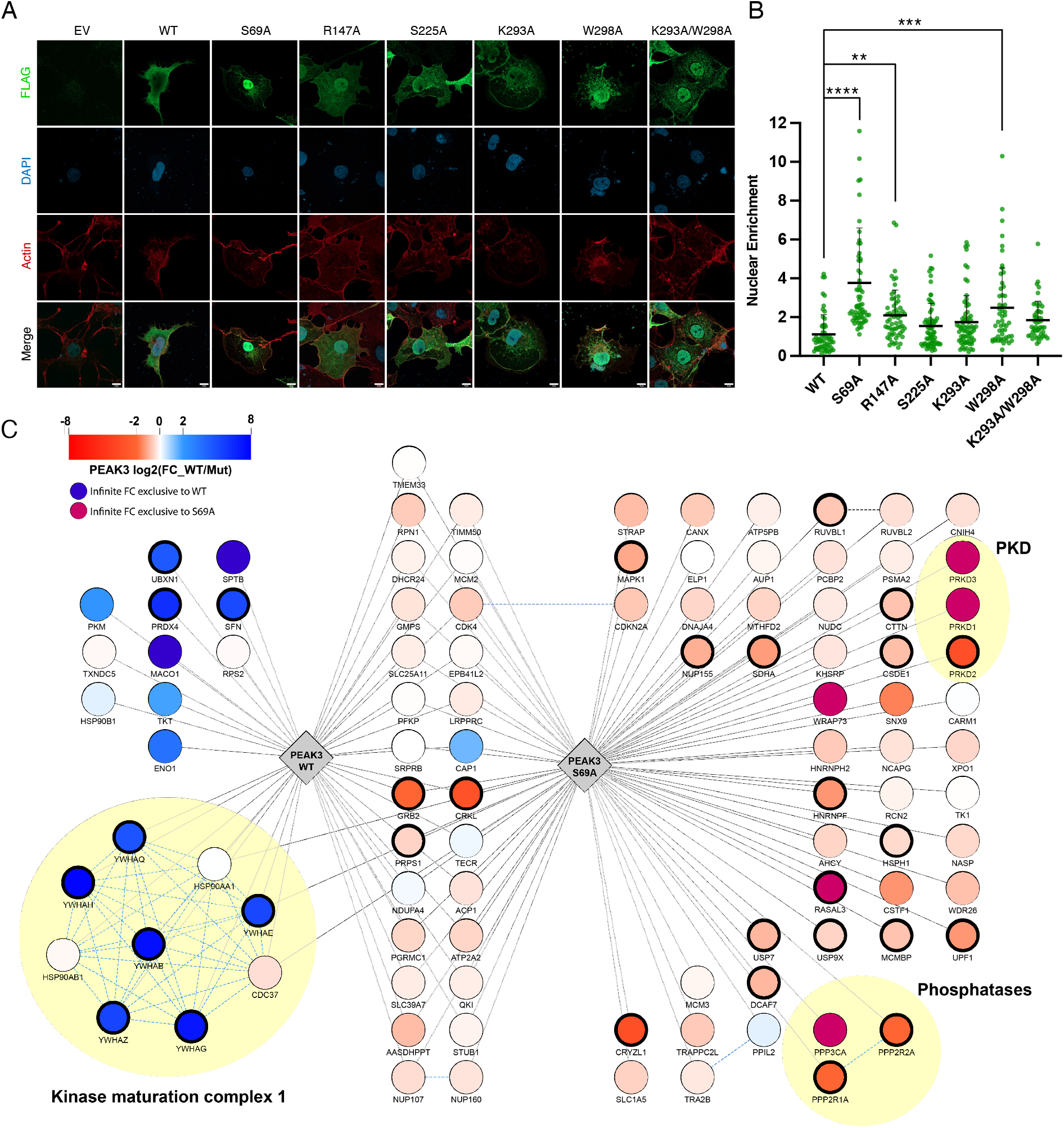
14-3-3 influences cellular localization and scaffolding range of PEAK3. A) Immunofluorescence-based imaging of FLAG-tagged PEAK3 transiently expressed in COS-7 cells. Representative confocal microscopy images show cells transfected with an empty vector (EV) control or PEAK3 variants: WT, S69A, R147A, S225A, K293A, W298A, or K293A/W298A. PEAK3 was detected with an anti-FLAG antibody (green), and cells were further stained with DAPI (blue, nucleus) and iFluor-647 conjugated phalloidin (red, actin). Scale bars: 10 μm. B) Quantification of relative nuclear enrichment of PEAK3 under conditions of impaired 14-3-3 binding. The ratio of fluorescence intensity in the green channel after background subtraction measured in the nucleus to the fluorescence intensity of the non-nuclear portion of the cell is plotted for each PEAK3 variant; see Materials and Methods for details. Data are plotted as the mean with standard deviation, combining all cells from at least 3 independent experiments (n = 57, 65, 60, 69, 71, 53, and 48 total cells for WT, S69A, R147A, S225A, K293A, W298A, and the K293A/W293A, respectively), *p < 0.05. C) Spectrum of protein-protein interactions engaged by the WT or S69A PEAK3 analyzed by immunoprecipitation/mass spectrometry approach. Edges represent significant specific interactions of PEAK3 WT or S69A as determined by SAINTexpress (BFDR < 0.05) compared to negative control (non-transfected cells). Node colors represent the log2 fold change enrichment determined by MSstat between WT and S69A PEAK3 (bold node border represents significant adjusted p value < 0.05). CORUM complexes are indicated by yellow circles.

### Loss of 14-3-3 binding and nuclear redistribution of PEAK3 enables novel protein-protein interactions

The redistribution of PEAK3 to the nucleus in the absence of 14-3-3 binding prompted us to examine if this change in localization modulates PEAK3’s function as a molecular scaffold. We used immunoprecipitation coupled with mass spectrometry to map the binding partners of WT PEAK3 and the 14-3-3 binding-deficient S69A mutant transiently expressed in HEK293 cells. Remarkably, we observed that while the WT PEAK3 predominantly interacted with 14-3-3 proteins and several motility factors, as previously described^12^, the S69A mutant showed a more diverse set of binding partners, with some interactors being present in both conditions but significantly increased in the absence of 14-3-3 binding **(Figure 6C)**. Notable changes included an increase in Grb2 and CrkL binding to the S69A mutant compared to WT PEAK3. Some interactions were only observed in the absence of 14-3-3 binding, including binding of PEAK3 to a type 2A serine/threonine phosphatase complex composed of PPP2R1A, PPP2R2A, and PPP3CA, and to three protein kinase D (PKD) members, PRKD1, PRKD2, and PRKD3. This substantial expansion of the PEAK3 interactome in the absence of 14-3-3 binding strongly suggests that one of the primary functions of 14-3-3 binding to PEAK3 is to regulate its scaffolding potential, either by directly blocking interactions between PEAK3 and its binding partners or by restricting PEAK3 shuttling between the cytosol and the nucleus.

The enhanced interaction of PKD kinases with the 14-3-3 binding-deficient mutant of PEAK3 suggests that these kinases could mediate PEAK3 Ser 69 phosphorylation. Indeed, the sequence of PEAK3’s 14-3-3 binding site, which encompasses Ser 69, is consistent with a PKD substrate consensus site **(Figure 7A)**. Treatment of HEK293 cells with increasing concentrations of the PKD-specific inhibitor CRT0066101 dihydrochloride^33^ resulted in a dose-dependent decrease of endogenous 14-3-3 co-immunoprecipitation with FLAG-tagged WT PEAK3, relative to that of DMSO treated cells **(Figure 7B)**. Thus, PKD kinases are likely candidates for regulation of the PEAK3/14-3-3 interaction, and hence PEAK3 scaffolding potential more broadly.

**Figure 7:**
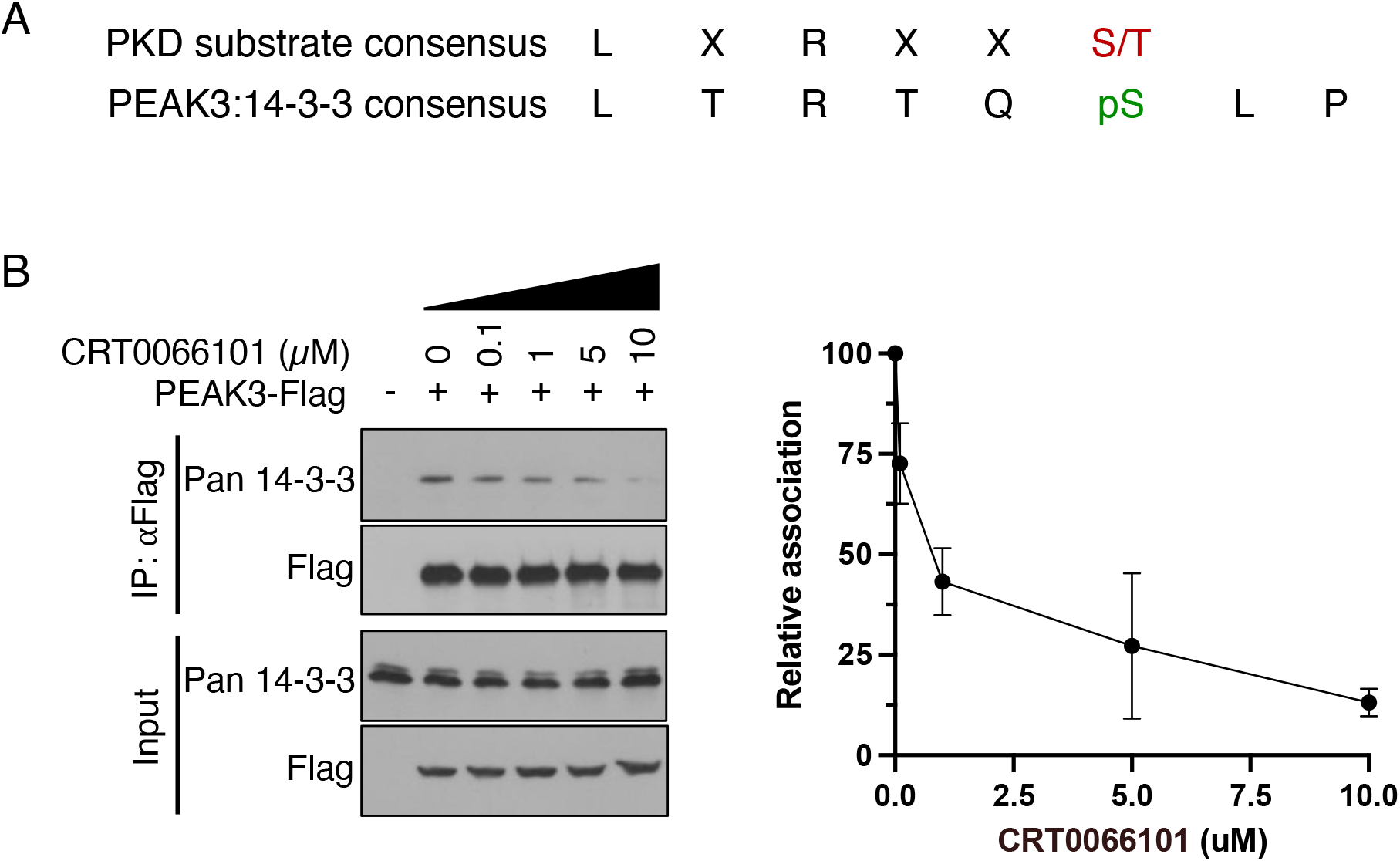
PKD regulates the PEAK3/14-3-3 interaction. A) Consensus sequence of PKD substrates aligned with PEAK3’s 14-3-3 consensus site. B) Left: Representative co-immunoprecipitation of endogenous 14-3-3 with FLAG-tagged WT PEAK3 transiently expressed in HEK293 cells treated with DMSO or increasing concentrations of the PKD inhibitor CRT0066101 dihydrochloride. Protein levels were detected with the indicated antibodies by Western Blot. Right: Quantification of co-immunoprecipitation data representative of 3 independent experiments.

## Discussion

Here, we present the first structure of the PEAK3 pseudokinase and the first structural characterization of any full-length PEAK family member. The N-terminal domain of PEAK3 in our structure is largely disordered, but its presence was critical for capturing PEAK3 in complex with the endogenous 14-3-3 dimer. 14-3-3 proteins have been consistently identified in unbiased mass spectrometry studies as top PEAK3 interaction partners^12-14^. In our experiments, this interaction was key to purifying full-length PEAK3 and may have played a stabilizing role yielding a higher resolution cryo-EM reconstruction **(SI Figure 5)**. While several putative 14-3-3 binding motifs can also be found in the N-terminal domains of PEAK1 and PEAK2 **(SI Figure 2B, 2C)**, it is unclear whether 14-3-3 proteins play a similar role for these family members, since the use of N-terminal truncated constructs in previous structural studies of PEAK1 and PEAK2 eliminated the chances of their isolation in complex with 14-3-3s^11,16,17^.

In the PEAK3/14-3-3 structure, two distinct interfaces contribute to complex formation. The first, canonical interface, involves binding between the 14-3-3 binding motifs located in the N-terminal domains of each PEAK3 monomer and the conserved 14-3-3 amphipathic grooves. The second interface is unique and contrasts with commonly observed secondary interactions, where the 14-3-3 client sit within or on top of the 14-3-3 cradle and typically interacts with both 14-3-3 dimer molecules. Instead, the secondary interface in the PEAK3/14-3-3 complex involves only one monomer of 14-3-3, which forms an extensive interface with the PEAK3 dimer. Presumably, increased dynamics associated with such architectures might have prevented structure determination in other complexes. Interestingly, solution studies designed to characterize dynamic protein-protein interactions, such as small angle X-Ray scattering (SAXS), have previously pointed to similarly asymmetric binding poses between 14-3-3 and the CaMKK2 and DAPK2 kinases^34,35^. While in our experiments 14-3-3 binding clearly reduces the overall diversity of PEAK3’s interactome, the unique architecture of the PEAK3/14-3-3 complex captured in our structure leaves the unstructured PEAK3 N-termini accessible. As these contain numerous consensus binding sites for key signaling proteins, having these regions exposed may play an important role in tuning of PEAK3 scaffolding function and/or initial recruitment of PEAK3 complexes.

Our structure highlights the conservation of SHED-dependent dimerization across all PEAK pseudokinases, as previously anticipated based on structural models^12^. We propose here why dimerization of PEAKs is essential for binding of its signaling partners, a phenomenon consistently observed across studies^11,16,17^. Based on our finding that the PEAK3 SHED domain forms essential contacts with 14-3-3, largely contributing to the secondary interface of the PEAK3/14-3-3 interaction, we speculate that this binding mode might represent a universal model for how PEAK pseudokinases interact with other signaling molecules which depend on PEAK dimerization for binding. This hypothesis is supported by our observation that when 14-3-3 binding to PEAK3 is disrupted, exposing the SHED domain interface, the spectrum of PEAK3 interactions with other binding partners increases significantly.

The PEAK3 structure highlights unique structural features of its pseudokinase domain that are distinct from PEAK1 and PEAK2. The overall architecture of the PEAK3 pseudokinase homodimer within the PEAK3/14-3-3 complex, or alone solved at lower resolution, shows significant twisting between two pseudokinase domains instigated by rotations of the SHED domain helices, αJ and αL, relative to PEAK1 and PEAK2. These differences are thus independent from 14-3-3 binding and intrinsic to PEAK3. Within the putative ATP-binding site, PEAK3 resembles its paralogs, as it also has a pocket highly occluded by the side chains of multiple residues which engage the catalytic lysine. However, unlike in structures of PEAK1 and PEAK2, in which the αC helix and activation loop were either partially or not at all resolved, these structural elements are found intact in the PEAK3 structure due to their stabilization by the interaction with 14-3-3ε. The proline-rich sequence of the PEAK3 activation loop, which is positioned closest to 14-3-3ε, is distinct in PEAK3 compared to PEAK1 and PEAK2. This sequence is reminiscent of proline-rich SH3 domain-binding sites, suggesting a potential role for the PEAK3 activation loop in the recruitment or stabilization of SH3 domain-containing interaction partners. A number of characterized PEAK3 binding partners, such as CrkII and Grb2, contain two SH3 domains. The canonical proline-rich motifs found within PEAK3’s N-terminal domain may engage one of the SH3 domains to initiate the interaction with these partners, while the less conserved motif within the activation loop may act to further stabilize the complex, similar to what is seen in the PEAK3/14-3-3 complex.

The strong association between PEAK3 and 14-3-3 underscores a functional role for this interaction. 14-3-3s regulate their client proteins through several mechanisms, including inducing structural changes, protecting phosphorylation sites, altering their subcellular localization, and facilitating or blocking protein-protein interactions^19-22^. We show that 14-3-3 is not essential for PEAK3 dimerization, but rather controls subcellular localization of PEAK3 and prevents it from nuclear translocation. Since PEAK3 does not have its own nuclear import signal, it could be that 14-3-3 blocks PEAK3 from interacting with another protein that is responsible for delivery of PEAK3 to the nucleus. Our mass spectrometry analysis suggests that when freed from 14-3-3 binding, PEAK3 serves as a multi-protein scaffold, as its interactome is greatly expanded when 14-3-3 is not bound. Thus, 14-3-3 might exert a dual control over PEAK3 function: over its localization and extent of its protein-protein interactions.

Among notable PEAK3 interactors enriched in the absence of 14-3-3 binding are a PP2A serine/threonine phosphatase complex and the PKD family of serine/threonine kinases. PKDs play numerous important roles in the cell, including regulating secretion and vesicle transport through the trans-Golgi network, inducing migration and differentiation, attenuating JNK activation in response to EGFR stimulation, and stimulating the Ras and ERK pathways^36-41^. One way PKDs exert their pleiotropic effect is by phosphorylating 14-3-3 consensus motifs of their substrates, thereby promoting 14-3-3 binding and cytoplasmic sequestration of their substrates^42^. For example, RIN1, which in its unphosphorylated form inhibits the Ras/Raf interaction by associating with membrane-bound activated Ras, loses this ability upon PKD-dependent phosphorylation of its 14-3-3 consensus that promotes 14-3-3 binding and blocks RIN1 association with the membrane^43^. Another example is class IIa histone deacetylases, which can shuttle between the nucleus and cytoplasm and form complexes with transcriptional repressors in the nucleus; however, PKD phosphorylation enables 14-3-3 binding, restricting nuclear import of class IIa histone deacetylases ^44,45^. We show that phosphorylation of the 14-3-3 consensus site in PEAK3 is also PKD-dependent, suggesting that PKD might regulate the extent of PEAK3’s role as a scaffold via 14-3-3 binding. PP2A phosphatases, which we also found recruited to PEAK3, might oppose the PKD-mediated effects, constituting the opposing arm of this regulation.

How and if 14-3-3 regulates interactions of PEAK3 with other signaling proteins, such as MAPK1 and Grb2, is a topic of future investigation. PEAK3 overexpression and knockdown has been shown to increase and decrease MAPK1 phosphorylation in acute myeloid leukemia (AML) cells, respectively^14^, and 14-3-3 may play an important role in this regulation. Grb2 is known to interact with PEAK3 via the N-terminal domain Tyr 24 and has been shown to regulate cell invasiveness in breast epithelial cells^13^. The enriched interaction between Grb2 and PEAK3 in the absence of 14-3-3 binding reported here suggests that 14-3-3 might negatively regulate the oncogenic Grb2/PEAK3 signaling axis.

Our structure provides the first characterization of how the SHED domain supports the scaffolding functions of PEAK pseudokinases. It also underscores the remarkable conservation of pseudokinase architecture and dimeric organization across all PEAK pseudokinases, suggesting that all of them might utilize a similar interface for engaging with their signaling partners. At the same time, the unique features of the PEAK3 pseudokinase domain offer a possibility for conceptualizing paralog-specific therapeutics^12,13,18^. An expanding body of evidence demonstrating association of PEAKs with progression of human cancers makes them emerging targets of interest^3^. Binding of PEAK3 to 14-3-3 may play an important role in buffering the population of PEAK3 that is available to engage with its scaffolding substrates and thereby modulate the oncogenic potential of PEAK pseudokinases.

## Author Contributions

M.L and D.D. cloned, expressed, and optimized the purification of the PEAK3/14-3-3 complex and D.D. performed initial NS-EM analysis. M.D.P. expressed and H.T purified the PEAK3/14-3-3 complex for cryo-EM studies. H.T. optimized sample preparation for cryo-EM, collected data and built/refined structural models under the guidance of K.A.V. H.T. and M.L. generated constructs for immunoprecipitation and immunofluorescence studies, performed immunoprecipitation studies and H.T. performed data analysis. M.D.P. performed immunofluorescence studies and analyzed results. A.F. performed mass spectrometry experiments and analyzed the data, under the supervision of N.J.K. K.P performed analysis of PEAK3 sequence conservation. Figures were generated by H.T, M.D.P., A.F., and K.P. under the guidance of N.J. and K.A.V and the manuscript was written by H.T., M.D.P., N.J., and K.A.V, with edits from M.L., D.D, and K.P. and with approval from all authors. The project was supervised by N.J. and K.A.V.

## Acknowledgments

We thank D. Bulkley and the rest of the staff at the UCSF cryo-EM imaging facilities for help with cryo-EM data collection, and M. Gupta for providing n-Octyl-Π-D-glucoside. We thank the members of the Jura and Verba labs for helpful discussions. This work was supported by the UCSF Program for Breakthrough Biomedical Research to K.V. and N.J., QBI Independent Research Fellowship to K.V., NIH/NIGMS R35-GM139636 to N.J., and the pilot project grant within the U54 CA209891 to N.J. and N.J.K.

## Materials and Methods

### Plasmids and cell culture

The human PEAK3 gene was synthesized by GenScript and subcloned into the pcDNA4/TO vector with a C-terminal 3xFLAG tag, 1xFLAG tag, or HA tag. Mutations were introduced using Quikchange mutagenesis (Agilent). All constructs were verified via DNA sequencing (Elim Biopharmaceutical). HEK293 and COS-7 cells were maintained at 37 ºC and 5% CO2 and cultured in Dulbecco’s modified Eagle media (Life Technologies) supplemented with 10% FBS (Hyclone) and 1% penicillin/streptomycin (Life Technologies). PEAK3 constructs were transiently transfected into cells 24 hours prior to imaging or cell lysis using Lipofectamine 3000 (Invitrogen) according to the manufacturer’s protocol.

### PEAK3 expression and purification

Expi293F suspension cells (Thermo Fisher Scientific Cat #A14527) were maintained at 37 ºC and 8% CO2 while rocking at 125 RPM (Thermo Fisher Scientific Cat #88881101 shaker, 125 ml non-baffled sterile cap vented flasks). The PEAK3-1xFLAG construct was transfected into 30 ml of Expi293F culture at a cell density of 4.0 × 10^6^ cells/ml using ExpiFectamine 293 transfection reagent (Thermo Fisher Scientific Cat #A14524) according to the manufacturer’s protocol at 1 μg DNA / 1 ml. Enhancer was added 18 hours post-transfection, and cells were collected 48 hours post-enhancer addition in an Allegra X-14 centrifuge (Beckman Coulter) at 1,000 xg for 10 min. Pellets were flash frozen in liquid nitrogen for later purification or resuspended in binding buffer (50 mM Tris-HCl, pH 7.5, 150 mM NaCl, 2 mM MgCl2, 2 mM DTT) with 1 mM NaF, 1 mM NaVO4, DNase I, and a cOmplete mini EDTA-free protease inhibitor cocktail tablet (Roche) supplemented fresh. If frozen, pellets were thawed on ice and homogenized by gentle pipetting. Cells were lysed using an EmulsiFlex-C5 homogenizer (Avestin) at 10,000–15,000 psi, and the cell lysate was spun down in an Avanti centrifuge with a JLA 25.50 rotor at 20,000xg for 40 min, at 4 °C. The clarified lysate was then incubated with 125 µL of G1 Flag resin (Genscript) per 30 ml culture overnight at 4 °C with rotating before washing with 50 bead volumes of lysis buffer. Resin was then incubated with 10 bead volumes of elution buffer (lysis buffer + 0.25 mg/ml FLAG peptide [SinoBiological]) for 1 hour at 4 °C with rotating and protein was eluted by gravity flow. The eluate was concentrated using an Amicon Ultra-0.5 30k MWCO centrifugal filter (Millipore), before being loaded and separated on an S200 10/300 GL column (GE Life Sciences). PEAK3/14-3-3 peak fractions were collected, concentrated to 20 µM and flash frozen.

### Sample preparation for cryo-electron microscopy imaging

Peak fractions of the PEAK3/14-3-3 complex were pooled and concentrated using an Amicon Ultra-0.5 30k MWCO centrifugal filter (Millipore). Immediately before preparing cryo-EM grids, a final concentration of 0.1% n-Octyl-β-D-glucoside (Affymetrix) was added to a 7 µM sample of the PEAK3/14-3-3 complex to limit orientation bias of PEAK3/14-3-3 complex particles. 3 µL of this sample was then applied to negatively glow discharged Quantifoil R1.2/1.3 300 mesh Au holey-carbon grids, blotted using a Vitrobot Mark IV (FEI), and plunge frozen in liquid ethane at 5 °C, 100% humidity, 4–8 s blot time, and 4 blot force.

Grids were imaged on a 300-keV Titan Krios (FEI) with a K3 direct electron detector (Gatan) and a BioQuantum energy filter (Gatan) using SerialEM v3.8.6 and Digital Micrograph v3.31.2359.0. Data for the PEAK3/14-3-3 complex were collected in super-resolution mode at a physical pixel size of 0.835 Å pixel^−1^ with a dose rate of 16.0 e^−^ pixel^−1^ s^−1^ (operated in non-CDS mode). Images were recorded with a 3 s exposure over 120 frames with a dose rate of 0.57 e^−^ Å^−2^ per frame.

### Image processing and 3D reconstruction

Raw movies were corrected for motion and radiation damage with MotionCor2^46^ during collection launched via Scipion^47^, and the resulting sums were imported in CryoSPARC2^48^. Micrographs were curated based on ice thickness, CTF fit resolution, and particle distribution, yielding a final stack of 80% of total micrographs collected. Micrograph contrast transfer function (CTF) parameters were estimated with the patch CTF job in CryoSPARC2. Particles were initially picked with blob picking, extracted, and 2D-classified yielding both PEAK3 only particles and PEAK3/14-3-3 complex particles. Particles were then template picked with low-pass filtered (to 20 Å) 2D class averages of the PEAK3/14-3-3 complex 2D class averages, and the resulting picks were extracted with 2× Fourier cropping and subjected to iterative rounds of ab initio and heterogeneous refinements (see processing workflow chart). Finally, unbinned particles were re-extracted and run through non-uniform refinements to achieve reconstruction with the highest resolution. The final reconstruction of PEAK3/14-3-3 complex used for model building included 169,563 particles with C1 symmetry and resulted in an overall resolution of 3.1 Å by gold standard FSC (GS-FSC) cut-off of 0.143 (masked), sharpened with -107.4 B-factor.

Particles in the same dataset were also template picked with low-pass filtered (to 20 Å) 2D class averages of the PEAK3 homodimer alone and processed similarly to PEAK3/14-3-3 complex particles. To alleviate streaking artifacts due to the orientation bias of PEAK3 homodimer particles in the reconstructions from CryoSPARC2, unbinned particles of the PEAK3 homodimer were re-extracted and imported into RELION3^49^ where they were further 3D-classified. The final reconstruction of the PEAK3/14-3-3 complex used for model building included 32,734 particles and resulted in an overall resolution of 4.9 Å by GS-FSC (masked).

Each map was assessed for local and directional resolutions through ResMap^50^ and 3DFSC^51^ server, respectively. For the PEAK3/14-3-3 complex reconstruction, the pseudokinase domain of the monomer of PEAK3 not interfacing with the 14-3-3 dimer was at local resolutions between 3.5–5.5 Å, while the remainder of the complex was at local resolutions between 2.5–3.5 Å.

### Model building, refinement, and validation

Model building for both the PEAK3/14-3-3 complex and the PEAK3 homodimer began with fitting of the AlphaFold2 models of PEAK3, with the N-terminal extensions truncated due to low confidence scores, and 14-3-3 isoforms ε and β into the corresponding 3D volumes in ChimeraX^52^. The PEAK3/14-3-3 complex and PEAK3 homodimer models were fit into the respective cryo-EM maps with FastRelax protocol in Rosetta in torsion space^53^. For the PEAK3/14-3-3 complex, the PEAK3 N-terminal peptides within the 14-3-3 cradle were built manually in Coot and the full complex model was further manually examined and refined in Coot and ISOLDE^54,55^. Per-atom *B*-factors were assigned in Rosetta indicating the local quality of the map around that atom^53^. MolProbity, as part of the Phenix validation tools, was used to assess quality of the models^56,57^. The residues modeled for the PEAK3/14-3-3 complex are as follows: PEAK3-Chain A (128-406, 419-473), PEAK3-Chain B (130-406, 415-472), 14-3-3β (3-232), 14-3-3ε (3-233), PEAK3 N-terminus-Chain E (66-73), and PEAK3 N-terminus-Chain P (66-72). Residues modeled for the PEAK3 homodimer are PEAK3-Chain A (128-406, 419-473) and PEAK3-Chain B (130-406, 415-472). Q-scores were calculated using the appropriate plugin in UCSF Chimera^58,59^.

### Identification and structural alignment of 14-3-3/client complexes

A search was carried out for 14-3-3/client complex structures in the Protein Data Bank using the Advanced Search Query Builder via the sequence similarity feature. Structures with a sequence similarity of 30% to 14-3-3ε and an E-value cutoff of 0.1 were manually curated, removing 14-3-3 structures that only contained client peptides, with no globular domains present. Following curation, structures were aligned in ChimeraX^52^ on the 14-3-3 dimer to visualize different binding modes. Structures exhibiting binding mode 1 include: PDB IDs: 2O98, 5N6N, 6UAN, 6Q0K, 7MFE, 7MFF, 1IB1, 6XAG, 6U2H. Structures exhibiting binding mode 2 include: PDB IDs: 3AXY, 6GN0, 6GNK, 6GNJ, 6GN8, 5LTW, 6GNN.

### Co-immunoprecipitation and Western blot analysis

HEK293 cells were seeded on 100 mm dishes at a density of 5.0 × 10^6^ cells/dish and transfected the following day. 24 hours post transfection, cells were washed two times on ice with ice cold PBS followed by lysis (0.5% Triton X-100, 0.5% NP-40, 150 mM NaCl, 50 mM Tris pH8.0, 1 mM NaF, 1 mM Na_3_VO_4_, 1 mM EDTA, cOmplete mini EDTA-free protease inhibitor cocktail [Roche]). Cells were scraped and incubated with rotation at 4 °C for 45 min to ensure complete lysis. Lysates were clarified by centrifugation for 10 min at 15,000 rpm at 4 °C. The clarified lysates were pre-cleared with 100 μL Protein A beads (Novex) for 30 min at 4 °C with rotation. The pre-cleared lysates were then incubated with 100 µL of antibody/protein A complexed beads (anti-FLAG [mouse, Sigma, F1804] or anti-HA [mouse, SCBT, sc-7392]) per 100 mm plate overnight at 4 °C with rotation. The beads were washed three times with lysis buffer. The bound proteins were eluted from the beads using SDS-loading buffer and were boiled at 95 °C for 10 min prior to SDS-PAGE and analysis by Western blot. Samples were run on 4-15% acrylamide gels and transferred onto PVDF membranes. Membranes were blocked in 5% milk in TBS and 0.1% Tween-20 (TBS-T) for 30 min at room temperature, followed by incubation with primary antibodies diluted in blocking buffer (anti-FLAG [rabbit, CST, 2368S], and anti-HA [mouse, SCBT, sc-7392]) at 4 °C overnight. Next, blots were incubated with secondary antibodies diluted in blocking buffer (anti-IgG Veriblot, Abcam, ab131366) for 1 hour at room temperature. ECL Western blotting detection reagent (GE) or ECL prime (VWR) were used for detection.

Western blot quantification was performed in FIJI Version 2.0.0 using the gel analysis tool. First, the 14-3-3 IP signal was normalized against the corresponding IP FLAG signal. Next, each 14-3-3 IP signal was normalized against the WT 14-3-3 (secondary interface IP) or DMSO control (PKD inhibition IP) signal. Statistical significance was determined using One-way ANOVA Dunnett’s multiple comparisons test in Prism 9 (Graphpad Software, Inc.).

### Immunofluorescence

COS-7 cells (1.0 × 10^5^ per well) were plated onto glass coverslips or in glass bottom petri dishes (MatTek, P35GCOL-1.5-14-C) and transfected the following day. 24 hours post transfection, cells were rinsed twice with PBS, followed by fixation in 3.7% formaldehyde for 10 min at room temperature. Cells were then rinsed three times with PBS, followed by blocking and permeabilization with the blocking/permeabilization buffer (PBS, 2.5% BSA, 0.1% Triton X-100) for 20 min at room temperature. Next, the cells were incubated with primary antibodies (anti-FLAG [rabbit, CST, 2368S]) diluted in the blocking/ permeabilization buffer for 1 hour at room temperature. The cells were then rinsed three times with PBS, followed by incubation with secondary antibodies (AlexaFluor 488 anti-rabbit IgG [donkey, Life Technologies, A21206]) diluted in blocking buffer for 1 hour at room temperature. During the last 20 min of this incubation, iFluor-647-Phalloidin (Cayman Chemical Company, 20555) was added at a 1:1000 dilution to stain actin. Coverslips were mounted onto glass coverslips with Prolong Gold AntiFade with DAPI (Cell Signaling Technologies) to stain the nucleus. Images were acquired on a Zeiss LSM 800 confocal laser scanning microscope using a 63x oil objective.

Image analysis was performed in FIJI. The nucleus and total cell were manually outlined using the nuclear (DAPI channel) and actin (AF647 channel) stains, respectively, and then the fluorescence intensity of PEAK3 (AF488 channel) in each region was measured. Nuclear enrichment in each cell was determined post background correction as the ratio of average intensity in the nucleus to average intensity of the whole cell, excluding the nucleus. Each condition was imaged in at least three independent experiments, and the data from all cells across all experiments were combined. Statistical significance was determined using One-way ANOVA Dunnett’s multiple comparisons test in PRISM (Graphpad Software, Inc.).

### Immunoprecipitation/Mass spectrometry

#### Immunoprecipitation of PEAK3

One 100 mm plate of cells (∼80% confluency) was washed with ice-cold Dulbecco’s PBS and lysed with 500 μl of ice-cold lysis buffer containing 100 mM KCl, 5 mM MgCl2, 20 mM Tris-HCl pH 8, 0.1% Tween-20, 0.1% NP40, 10% glycerol, 1× protease inhibitor cocktail (Roche, complete mini-EDTA free), and 1× phosphatase inhibitor cocktail (Roche, PhosStop). Lysates were sonicated and then clarified by centrifugation at 13,000 xg for 15 minutes at 4°C. For FLAG purification, 25 μl of bead slurry (Anti-Flag M2 Magnetic Beads; Sigma-Aldrich) was washed twice with 1 ml of ice-cold wash buffer (100 mM KCl, 5m M MgCl2, 20 mM Tris-HCl pH 8, 0.1% Tween-20, 0.1% NP40, 10% glycerol), and 2 mg lysate was incubated with the anti-FLAG beads at 4 °C with rotation for 3 hours. After incubation, flow-through was removed and beads were washed 3x with 1 ml of wash buffer and twice with 1 ml reduced wash buffer (100 mM KCl, 5m M MgCl2, 20 mM Tris-HCl pH 8). Bound proteins were eluted by incubating beads with 15 μl of 100 ug/ml 3xFLAG peptide in 0.05% RapiGest in wash buffer for 15 min at room temperature with shaking. Supernatants were removed and elution was repeated.

#### Preparation of samples for MS

For immunoprecipitated PEAK3 from 293T, eluates were combined and 10 μl of 8 M urea, 250 mM Tris, 5 mM DTT (final concentration ∼1.7 M urea, 50 mM Tris, and 1 mM DTT) was added to give a final total volume of ∼45 μl. Samples were incubated at 60 °C for 15 min and allowed to cool to room temperature. IODO was added to a final concentration of 3 mM and the mixture was incubated at room temperature for 45 min in the dark. DTT was added to a final concentration of 3 mM before adding 1 μg of sequencing-grade trypsin (Promega) and incubating at 37 °C overnight. Samples were acidified to 1% trifluoroacetic acid (TFA, pH < 2) with 20% TFA stock and incubated for 30 min before desalting on C18 stage tip. For purified PEAK3/14-3-3 complexes, 5ug of the purified protein complex was resuspended in 10 μl of 8 M urea, 50 mM Tris, 1 mM DTT. Samples were incubated at 60 °C for 15 min and allowed to cool to room temperature. IODO was added to a final concentration of 3 mM and the mixture was incubated at room temperature for 45 min in the dark. DTT was added to a final concentration of 3 mM and the sample was then diluted with 90 μl of 50 mM Tris before adding 0.1 μg of sequencing-grade trypsin (Promega) and incubating at 37 °C overnight. Samples were acidified to 1% trifluoroacetic acid (TFA, pH < 2) with 20% TFA stock and incubated for 30 min before desalting on C18 stage tip.

#### MS data acquisition and analysis

For AP-MS and purified PEAK3/14-3-3 complex experiments, samples were resuspended in 15 μl of MS loading buffer (1% formic acid) and 2 μl was separated by a reversed-phase gradient over a nanoflow 75 μm internal diameter × 25 cm long picotip column packed with 1.9 μM C18 particles (Dr. Maisch). Peptides were directly injected over the course of a 75 min acquisition into a Q-Exactive Plus mass spectrometer (ThermoFisher Scientific). Raw MS data were searched against the Uniprot canonical isoforms of the human proteome using the default settings in MaxQuant (version 1.6.12.0), with a match-between-runs enabled^60^. Peptides and proteins were filtered to 1% FDR in MaxQuant, and identified proteins were then subjected to PPI scoring. To quantify changes in interactions between WT and mutant baits, we used a label-free quantification approach in which statistical analysis was performed using MSstats^61^ from within the artMS Bioconductor R package. All raw data files and search results are available from the Pride partner ProteomeXchange repository under the PXD035574 identifier^62,63^. Detailed MS acquisition and MaxQuant search parameters are provided in the supplemental file.

#### PPI scoring

Protein spectral intensities as determined by MaxQuant search results were used for PPI confidence scoring by SAINTexpress (version 3.6.1) (37). For SAINTexpress, control samples in which bait protein was not transduced were used. Candidate PPIs were filtered to those that displayed a SAINTexpress BFDR <0.05.

#### Sequence bioinformatics

In order to generate the sequence logo for PEAK3’s 14-3-3 binding site, PEAK3 homologs were collected by BLAST search in the NCBI RefSeq protein sequence database. Orthologs of PEAK1 and PEAK2 were filtered out. Sequences thus collected were cleared of redundancy at identity threshold 0.8 using MMSEQ^64^ and aligned using Mafft^65^. The logo was generated using the Weblogo algorithm^66^.

