## Supplementary Material for "Structural insights into regulation of the PEAK3 pseudokinase scaffold by 14-3-3"

**Affiliations**

**Supplementary Figure 1.**

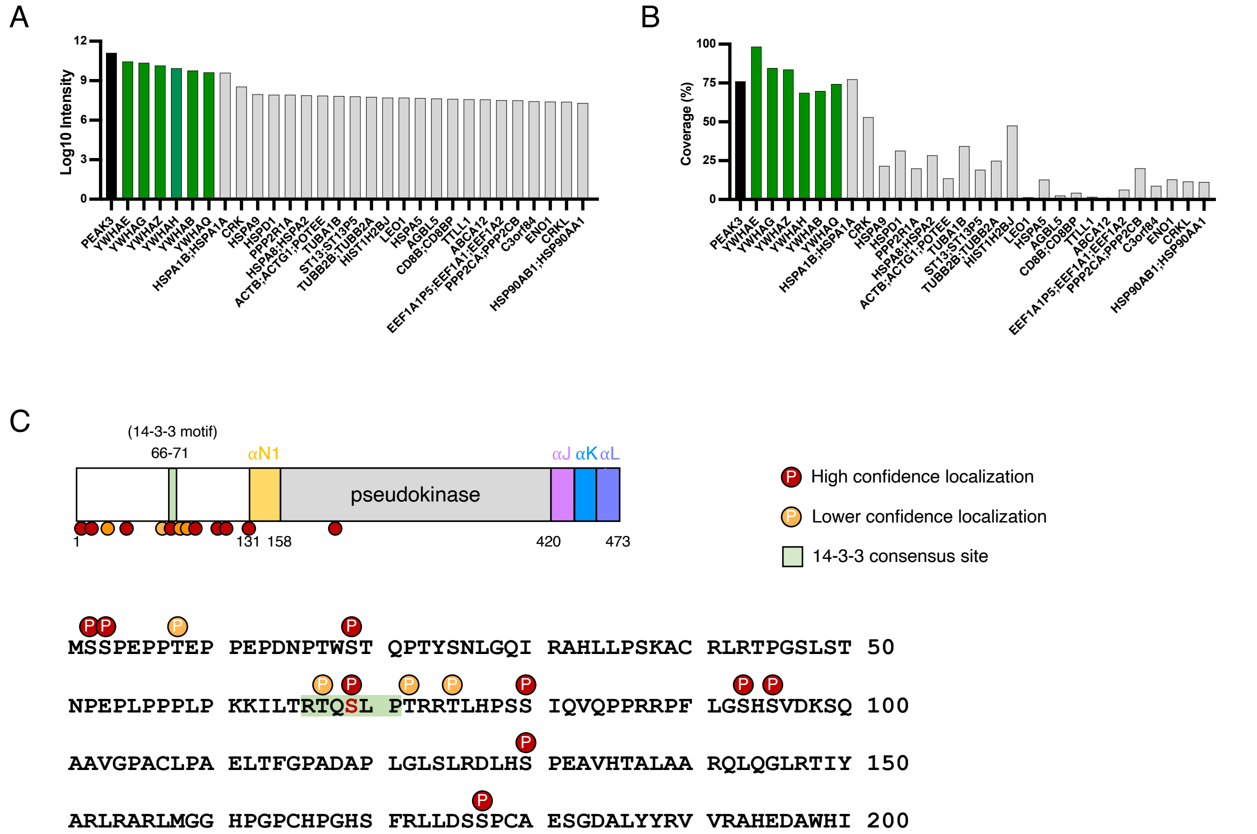


**SI Figure 1: Mass spectrometry analysis of the purified PEAK3/14-3-3 complex**

A) Bar chart representing the log10 protein spectral intensity for the top 30 proteins identified by MS of the purified PEAK3/14-3-3 complex, and B) their corresponding protein sequence coverage (black: PEAK3, green: 14-3-3, gray: other interactors). C) Schematic representation of the identified phospho-sites in PEAK3 by MS analysis (red: localization probability > 0.8; orange: localization probability < 0.8). The S69 phospho-site within the 14-3-3 consensus binding site (green) of PEAK3 represents a high confidence site.

### Supplementary Figure 2.

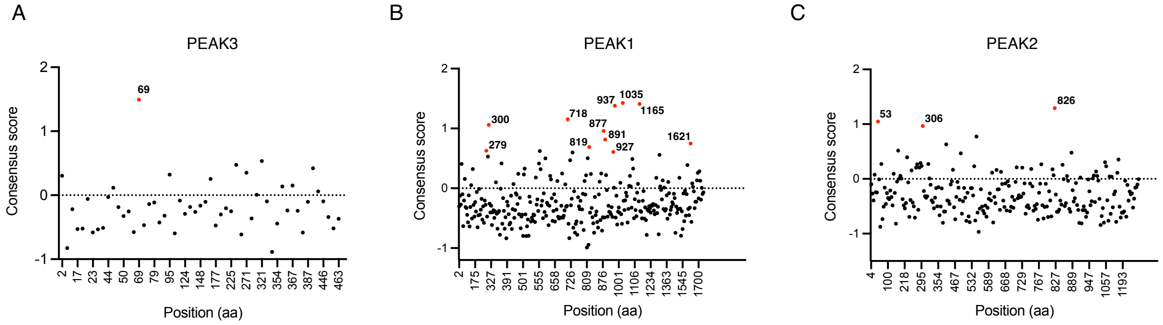


A

| Position | Sequence |
|---|---|
| 69 | ILTRTQ[S]LPTR |

B

| Position | Sequence |
|---|---|
| 279 | ANFRAN[T]LSPV |
| 300 | IPLRNK[S]LQRI |
| 718 | CLNRGQ[S]SPQR |
| 819 | TPVRPK[S]LFTS |
| 877 | PPPRST[S]SPYH |
| 891 | LLQRHF[T]NWTK |
| 927 | KPKRWI[S]FKSF |
| 937 | FFRRRK[T]DEED |
| 1035 | EKKRSH[S]SPSQ |
| 1165 | MIIRAN[T]EPIS |
| 1621 | LKEREY[T]RADL |

C

| Position | Sequence |
|---|---|
| 53 | PQPRAG[S]LPPP |
| 306 | QEKRGP[S]FPKE |
| 826 | IVSRAA[S]SPDG |

**SI Figure 2: Putative 14-3-3 binding sites in PEAK family members**

A-C) Sequence-based prediction of 14-3-3 consensus binding sites in PEAK family members using the 14-3-3-Pred webserver[25] based on three different classifiers (ANN, PSSM and SVM). Data points represent the amino acid position of the phosphorylated serine or threonine within the putative binding site. Sites, which score highly in all three prediction models, are colored red.

### Supplementary Figure 3.

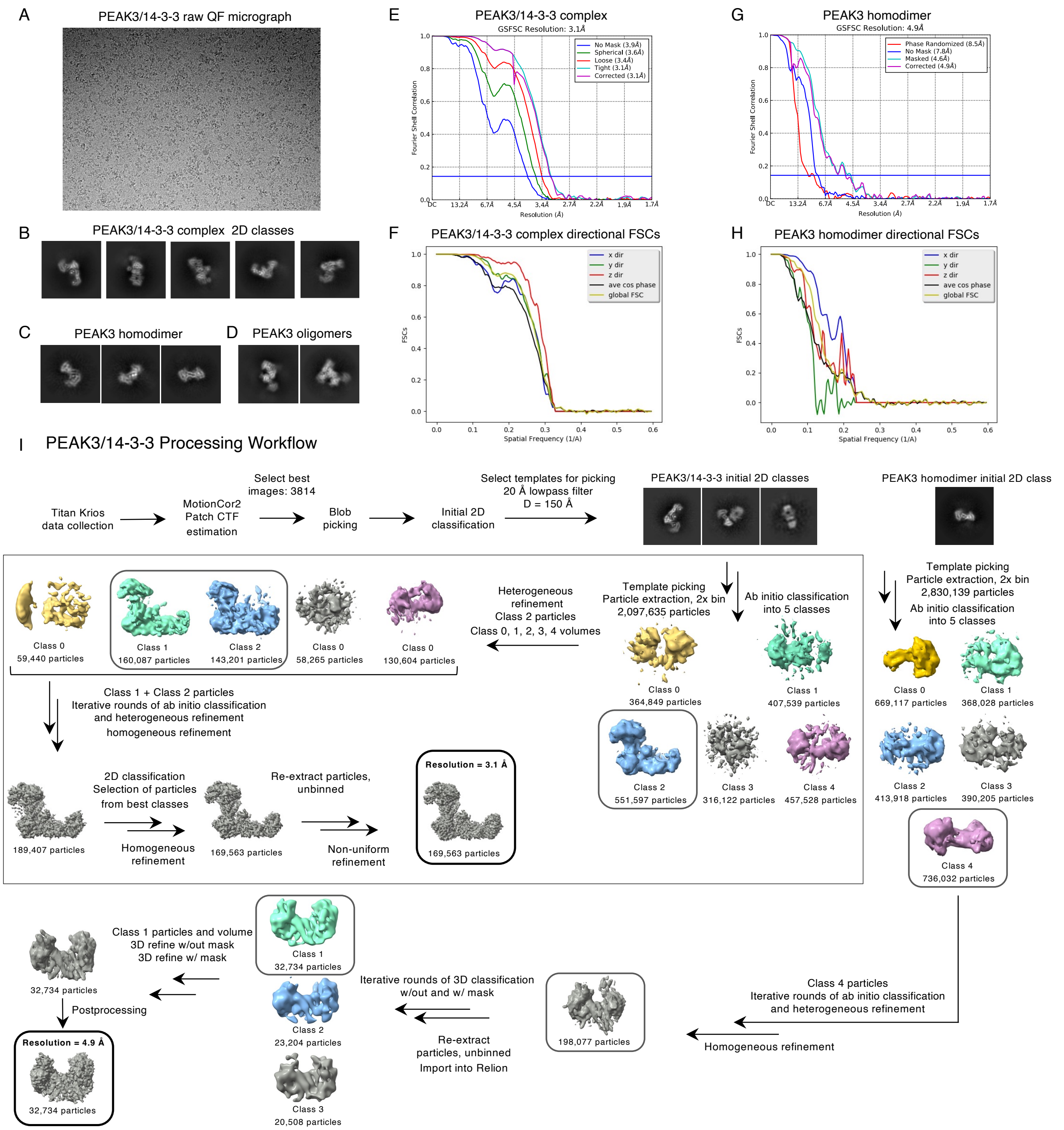


**SI Figure 3: Processing workflow, resolution estimation and map quality of PEAK3/14-3-3 complex dataset**

A) Representative micrograph of the PEAK3/14-3-3 complex sample on Quantifoil R1.2/1.3 300 mesh Au holey-carbon grids, from a dataset with 3814 micrographs. Example cryo-EM 2D class averages of particles corresponding to B) PEAK3/14-3-3 complex, C) PEAK3 homodimer, and D) PEAK3 oligomers in the absence or presence of 14-3-3 binding. Gold Standard Fourier Shell Correlation (GSFSC) of the final map used for model building of the E) PEAK3/14-3-3 complex from CryoSPARC2 with a reported resolution of 3.1 Å and final map for model building of the G) PEAK3 homodimer from RELION3 with a reported resolution of 4.9 Å. Directional FSCs of the F) PEAK3/14-3-3 and the H) PEAK3 homodimer calculated by 3DFSC server. I) Workflow for processing the PEAK3/14-3-3 complex dataset. Gray boxes indicate model and associated particle stack used for downstream processing. The final model is indicated with a bolded black box.

**Supplementary Figure 4.**

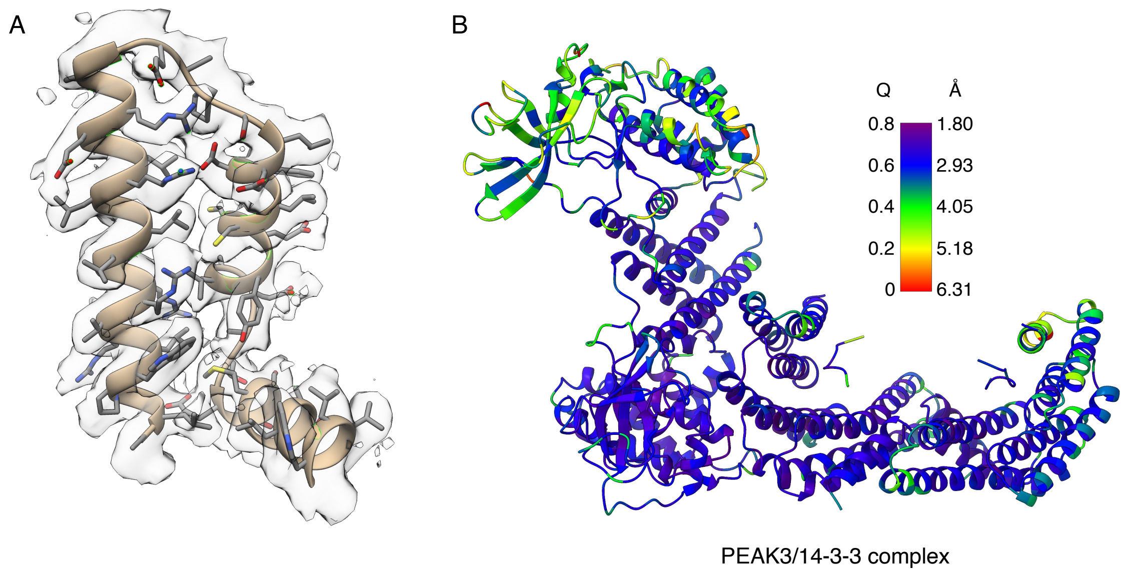


**SI Figure 4: Q-score analysis of PEAK3 cryo-EM structures**

A) Zoomed-in view of the cryo-EM map and corresponding model of the PEAK3/14-3-3 complex (residues 419-472) demonstrating features appropriate for reported resolution. B) PEAK3/14-3-3 complex model colored by estimated per residue Q-score. Color scale bar indicates corresponding estimated resolution in Å for reported Q-scores. Expected Q-score for a 3.1 Å structure is 0.569.

**Supplementary Figure 5.**

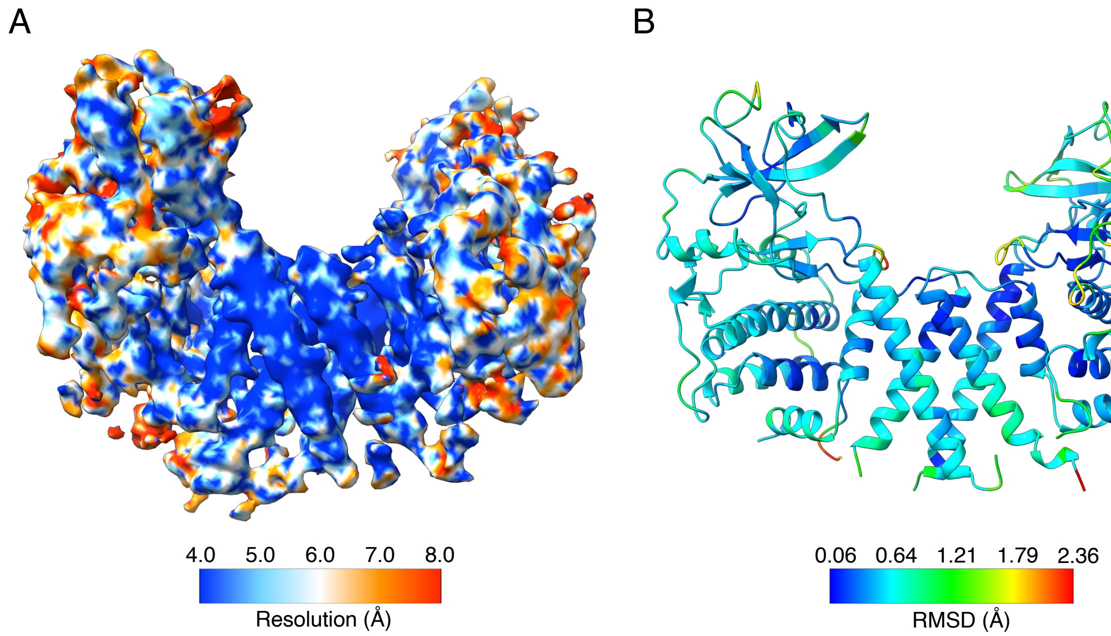


**SI Figure 5: Low-resolution structure of the PEAK3 homodimer**

(A) Cryo-EM map of PEAK3 homodimer colored according to local resolution determined by ResMap. B) Corresponding model of PEAK3 homodimer structure colored according by per residue RMSD (Å) relative to PEAK3 homodimer as part of the PEAK3/14-3-3 complex.

**Supplementary Figure 6.**

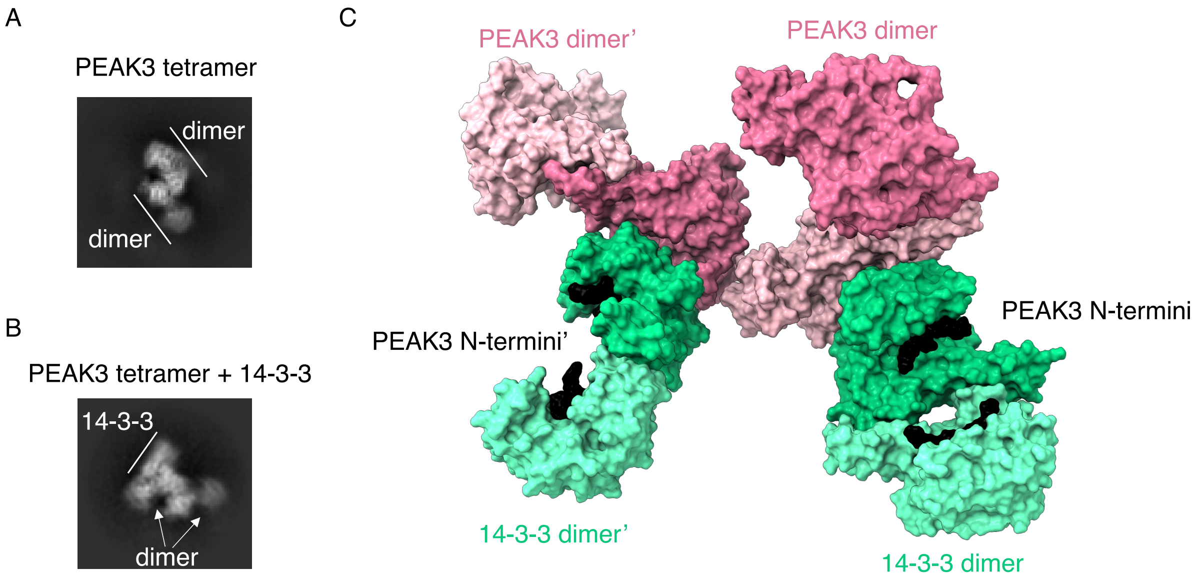


**SI Figure 6: PEAK3 oligomerization**

PEAK3 oligomerization in the A) absence and B) presence of 14-3-3 binding as demonstrated by cryo-EM 2D class averages. C) Putative oligomeric PEAK3/14-3-3 complex model derived from oligomeric interface identified from crystal packing of PEAK2 crystal structure (PDB ID: 5VE6). PEAK3 homodimer within the PEAK3/14-3-3 complex was first aligned with one of the two PEAK2 homodimers, followed by alignment of a second PEAK3 homodimer to the second PEAK2 homodimer.

**Supplementary Figure 7.**

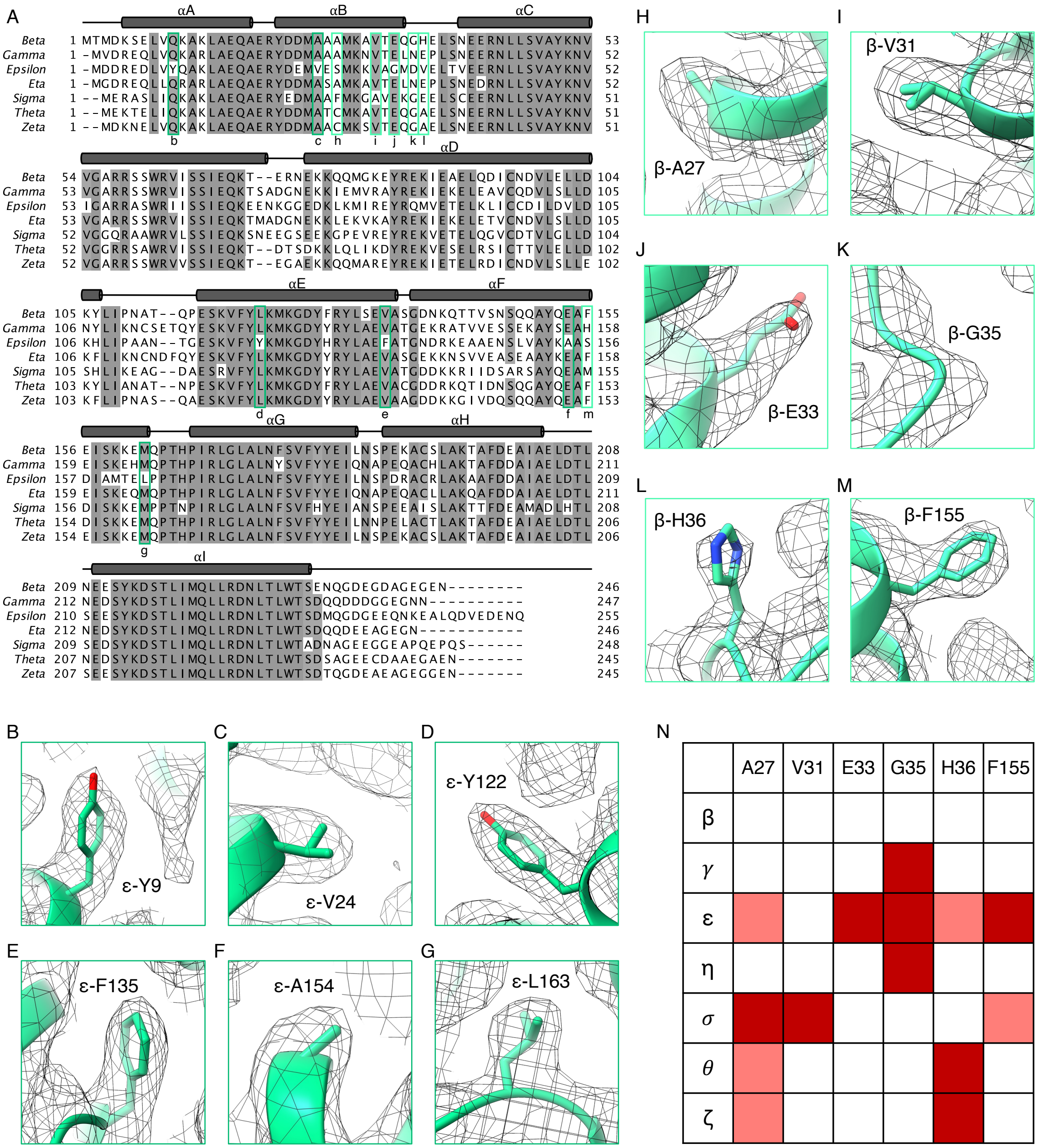


**SI Figure 7: Identification of 14-3-3 isoforms in PEAK3/14-3-3 complex**

A) Structure-based sequence alignment of human 14-3-3 isoforms depicting their secondary structure elements. Conserved residues are shaded in gray. B-M) Zoomed-in view of residue side chains in 14-3-3ε B-G) and in 14-3-3β H-M) overlayed with the cryo-EM map, which were used to determine the identity of the 14-3-3 monomers in the PEAK3/14-3-3 structure. N) Summary table of H-M residues and their compatibility with the cryo-EM map in all 14-3-3 isoforms. Dark red, light red and white squares indicate that the corresponding residue in a particular 14-3-3 isoform is not consistent, somewhat consistent or consistent with the cryo-EM density, respectively.

**SI Table 1. Cryo-EM collection, refinement and resulting model statistics.**

| | PEAK3/14-3-3 complex (EMD-27630) (PDB 8DP5) | PEAK3 homodimer (EMD-27684) (PDB 8DS6) |
|---|---|---|
| **Data collection and processing** | | |
| Magnification | 105,000x | 105,000x |
| Voltage (kV) | 300 | 300 |
| Total dose (e–/Å$^2$) | 69 | 69 |
| Dose rate (e-/physical pixel/sec) | 16 | 16 |
| Exposure per frame (sec) | 0.025 | 0.025 |
| Defocus range (μm) | -1.0 to -2.0 | -1.0 to -2.0 |
| Pixel size (Å) | 0.835 (physical) | 0.835 (physical) |
| Symmetry imposed | C1 | C1 |
| Initial particle images (no.) | 2097635 | 2608418 |
| Final particle images (no.) | 169563 | 32734 |
| Map resolution (Å)<br>FSC threshold (0.143) | 3.1 | 4.9 |
| Map resolution range (Å) | 2.5-5.5 | 4.0-8.0 |
| **Refinement** | | |
| Initial model used (PDB code) | AF- Q6ZS72<br>AF- P62258<br>AF- P31946 | AF- Q6ZS72 |

| | | |
|---|---|---|
| Model resolution (Å)<br>FSC threshold 0.5 (Masked) | 3.3 | 6.8 |
| Map sharpening *B* factor (Å$^2$) | -107.4 | -105 |
| Model composition<br>Non-hydrogen atoms<br>Protein residues | <br>8869<br>1145 | <br>5030<br>669 |
| *B* factors (Å$^2$)<br>Protein | <br>70.4 | <br>265.5 |
| R.M.S. deviations<br>Bond lengths (Å)<br>Bond angles (°) | <br>0.012<br>1.897 | <br>0.012<br>1.514 |
| Validation<br>MolProbity score<br>Clashscore<br>Poor rotamers (%) | <br>0.75<br>0.79<br>0.00 | <br>0.61<br>0.30<br>0.00 |
| Ramachandran plot<br>Favored (%)<br>Allowed (%)<br>Disallowed (%) | <br>98.93<br>1.07<br>0.00 | <br>98.64<br>1.36<br>0.00 |

102
